# Cyclin B3 implements timely vertebrate oocyte arrest for fertilization

**DOI:** 10.1101/2022.06.04.494806

**Authors:** Nora Bouftas, Lena Schneider, Marc Halder, Rebecca Demmig, Martina Baack, Damien Cladière, Melanie Walter, Hiba Al Abdallah, Camilla Kleinhempel, Janina Müller, Francesca Passarelli, Patrick Wehrle, Andreas Heim, Katja Wassmann, Thomas U. Mayer

**Affiliations:** Institut de Biologie Paris Seine, Sorbonne Université, 7 quai St. Bernard, 75252 Paris, France; CNRS UMR7622 Developmental Biology Lab, Sorbonne Université, 7 quai St. Bernard, 75252 Paris, France.; Department of Molecular Genetics, University of Konstanz, Universitätsstr. 10, 78464 Konstanz, Germany; Trenzyme GmbH, Byk-Gulden-Str. 2, 78467 Konstanz, Germany; German Cancer Research Center, INF 580, 69120 Heidelberg, Germany

## Abstract

To ensure successful offspring ploidy, vertebrate oocytes must halt the cell cycle in meiosis II until sperm entry. Emi2 is essential to keep oocytes arrested until fertilization. Yet, how this arrest is implemented exclusively in meiosis II and not prematurely in meiosis I remained enigmatic. Using mouse and frog oocytes, we show here that cyclin B3, an understudied B-type cyclin, is essential to keep Emi2 levels low in meiosis I. Direct phosphorylation of Emi2 at an evolutionarily highly conserved site by Cdk1/cyclin B3 targets Emi2 for degradation. In contrast, Cdk1/cyclin B1 is inefficient in Emi2 phosphorylation providing a molecular explanation for the requirement of different B-type cyclins for oocyte maturation. Cyclin B3 degradation at exit from meiosis I enables Emi2 accumulation and thus, timely arrest in meiosis II. Our findings illuminate the evolutionarily conserved mechanisms controlling oocyte arrest for fertilization at the correct cell cycle stage, essential for embryo viability.

## Introduction

The meiotic cell division is special, because two divisions – called meiosis I and II – have to be executed without intervening S-phase to generate haploid gametes (Bouftas and Wassmann, 2019; Petronczki et al., 2003). Additionally, the meiotic cell cycle program must be synchronized with the developmental program of gametogenesis such that key cell cycle events take place at the correct meiotic division (Lie et al., 2009; Ozturk, 2022). Examples are homologous chromosome segregation in vertebrates, that must take place only in meiosis I, and sister chromatid segregation that must take place only in meiosis II. Upon exit from meiosis I, but not meiosis II, S-phase must be suppressed and entry into the second meiotic division promoted (Petronczki *et al*., 2003). Vertebrate female meiosis poses an additional challenge with two cell cycle arrests that have to be implemented: A first arrest in prophase of meiosis I, required for immature oocytes to grow and accumulate maternal components for successful embryo development, and a second called CSF- (Cytostatic factor-) arrest in metaphase of meiosis II for mature oocytes to await fertilization (Masui and Markert, 1971; Ozturk, 2022). This second arrest is essential to prevent parthenogenetic divisions, i.e., embryo development without a paternal genome, and to maintain correct ploidy of the embryo.

Emi2 (also known as XErp1 in Xenopus) was identified as the key CSF component, that prevents parthenogenesis by directly inhibiting the APC/C (Liu et al., 2006; Madgwick et al., 2006; Schmidt et al., 2005; Shoji et al., 2006; Tung et al., 2005). Fertilization triggers the destruction of Emi2 resulting in APC/C activation and hence, meiotic exit (Hansen et al., 2006; Liu and Maller, 2005; Rauh et al., 2005; Suzuki et al., 2010). However, how CSF-arrest is synchronized with progression through the two meiotic divisions to ensure that oocytes arrest in meiosis II and not meiosis I, is largely unknown. The current model proposes two mechanisms to restrain Emi2 to meiosis II: dampened translation and degradation upon Cdk1/cyclin B-dependent phosphorylation (Natsumi Takei and Yamamoto, 2021; Ohe et al., 2007; Tang et al., 2008; Tung et al., 2007). But how Emi2 would be destabilized by Cdk1/cyclin B specifically in meiosis I, but not meiosis II, remained unclear.

Meiotic – like mitotic – cell cycle progression is driven by cyclin-dependent kinases (Cdks) in association with cell-stage specific cyclins (Morgan, 2007). Entry into and exit from M-phase is primarily driven by the synthesis and destruction of M-phase cyclins, i.e., cyclin A (cyclin A1 and A2) and cyclin B (cyclin B1 and B2). Switch-like activation of Cdk1 at M-phase entry is mediated by positive feedback loops between Cdk1 and its inhibitors and activators. Irreversible exit from M-phase results from cyclin B degradation upon ubiquitination by the E3 ligase APC/C (Anaphase-Promoting Complex/Cyclosome). A key open question is how the rapid activation of Cdk1 at M-phase entry is translated into a temporally precisely regulated sequence of phosphorylation events to ensure that the multitude of Cdk1 substrates is phosphorylated in the right order. This task becomes even more challenging for meiotic divisions, where exactly two waves of Cdk1 activation have to take place during two functionally distinct meiotic divisions. Differential substrate preference of the different Cdk1/cyclin complexes has been proposed as a mechanism underlying the timing of Cdk1 substrate phosphorylations (Hochegger et al., 2008). For example, homologue biorientation in Drosophila female meiosis I depends on the specific function of cyclin A2 (Bourouh et al., 2016). In mouse oocytes, cyclin A2 is important for sister chromatid segregation in meiosis II, and expression of a non-degradable mutant of cyclin A2 but not cyclin B1 during meiosis I leads to precocious sister chromatid separation, indicating that they target distinct substrates (Bouftas and Wassmann, 2019; Touati and Wassmann, 2016). But the most striking meiosis-specific role for a cyclin was recently discovered for the understudied B-type cyclin B3: Mutant mice harboring a genetic invalidation of *Ccnb3*, the gene coding for cyclin B3, are viable and males are fertile (Bouftas and Wassmann, 2019; Karasu et al., 2019; Karasu and Keeney, 2019; Li et al., 2019), indicating that cyclin B3 is not required in mitosis or male meiosis. Females, however, are sterile, with the vast majority of oocytes failing to enter meiosis II and instead remaining arrested in metaphase I. Cyclin B3 is degraded at exit from meiosis I (Karasu *et al*., 2019; Li *et al*., 2019), and translation is restrained to the first meiotic division, unlike cyclin B1 and B2 (Han et al., 2017). Yet, how cyclin B3 promotes anaphase I onset in oocytes, and why it is not required for second female meiosis as well as during male meiosis, has remained enigmatic.

Capitalizing on the amenability of mouse and frog oocytes to complementary experimental approaches, we show here that the key role of cyclin B3 is to ensure that CSF-arrest takes place only in meiosis II and not in meiosis I. Specifically, we demonstrate that Cdk1/cyclin B3 phosphorylates Emi2 at an evolutionarily highly conserved site in meiosis I, initiating a multisite phosphorylation cascade, which involves Plk1 and ultimately results in efficient degradation of Emi2. We prove that oocytes without cyclin B3 cannot complete the first meiotic division because they establish an Emi2-dependent cell cycle arrest at metaphase I. Notably, *in vitro* assays revealed that the phosphorylation site initiating Emi2 destruction is a very good substrate of Cdk1/cyclin B3, but not of Cdk1/cyclin B1. Furthermore, we show that cyclin B3 is degraded upon exit from meiosis I and does not re-accumulate. The absence of cyclin B3 from meiosis II is essential to install the CSF-arrest timely at metaphase II, and, accordingly, we confirm that untimely expression of cyclin B3 in meiosis II overrides the CSF-arrest. Hence, we discovered an evolutionarily conserved role for cyclin B3 in suppressing CSF activity in meiosis I. Cyclin B3 distinguishes itself from cyclin B1 in that it safeguards the generation of haploid oocytes competent to be fertilized, by ensuring they are arrested at the correct cell cycle stage, in meiosis II.

## Results

### Cyclin B3 prevents precocious CSF-arrest in mouse oocyte meiosis I

Homozygous *Ccnb3^−/−^* knock-out mouse oocytes arrest at metaphase I in a spindle assembly checkpoint-independent manner with stabilized securin and cyclin B (Figure S1) (Karasu *et al*., 2019; Li *et al*., 2019). Re-expression of wild-type (WT) mouse (*M.m.*) cyclin B3 but not of a kinase-dead hydrophobic patch mutant (MRL), rescued the metaphase I arrest (Karasu *et al*., 2019) demonstrating that Cdk1/cyclin B3 activity is essential for exit from meiosis I. Exogenously expressed cyclin B3 was degraded in an APC/C-dependent manner at exit from meiosis I and did not re-accumulate in meiosis II (Karasu *et al*., 2019).

The fact that cyclin B3 is essential for female but not male meiosis led us to hypothesize that cyclin B3 regulates a cell cycle transition specific to oocytes and absent in male meiosis. The phenotype of *Ccnb3^−/−^*oocytes is reminiscent of the metaphase II arrest of mature oocytes, so we wondered whether untimely CSF activity accounts for the observed metaphase I arrest. To test this hypothesis, *Ccnb3^−/−^* arrested oocytes were treated with strontium, an inducer of CSF release (O’Neill et al., 1991). Exit from meiosis I was examined by preparing chromosome spreads 1 hour after activation. Indeed, while all control-treated *Ccnb3^−/−^* oocytes displayed bivalents (connected pairs of homologous chromosomes), 74% of strontium-treated oocytes instead had segregated homologous chromosomes and showed dyads (pairs of sister chromatids), demonstrating successful progression through the first meiotic division (Figures 1A and S1). If Cdk1/cyclin B3 activity were required to suppress CSF-arrest in meiosis I, ectopic expression of cyclin B3 in meiosis II, when it is absent under physiological conditions (Han *et al*., 2017), should abrogate the CSF-arrest of mature mouse oocytes. Indeed, ectopic expression of WT, but not MRL mutant, *M.m.* cyclin B3 in mature CSF-arrested metaphase II wildtype oocytes resulted in rapid second polar body extrusion with separation of sister chromatids (Figures 1B and 1C), in accordance with (Meng et al., 2020). From these data we concluded that Cdk1/cyclin B3 negatively regulates CSF activity.

**Figure 1.**
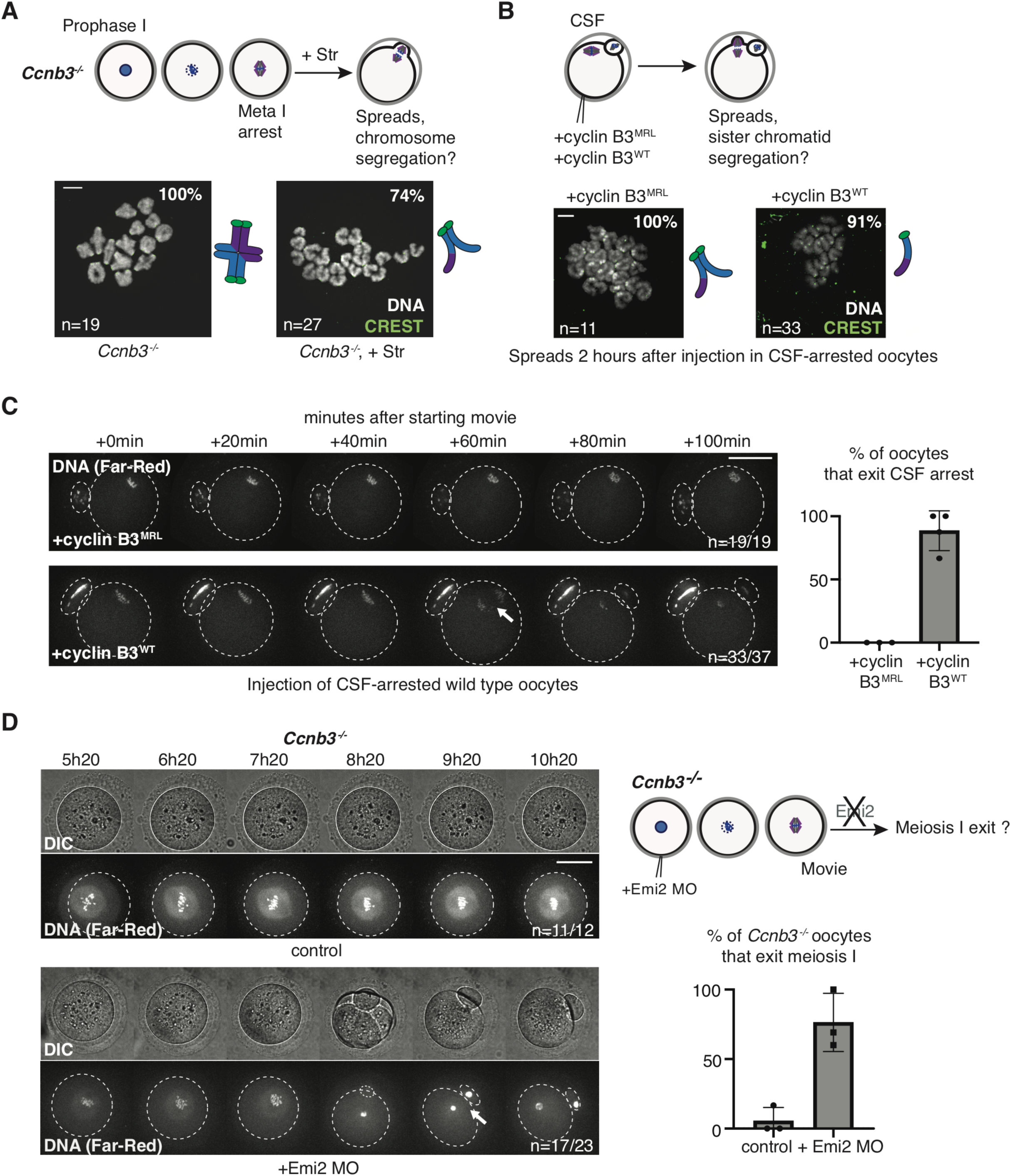
Cyclin B3 counteracts CSF-arrest in mouse oocytes. (A) Above, scheme of experimental setting to activate *Ccnb3^−/−^* metaphase I-arrested oocytes with strontium (+Str). Below, chromosome spreads of *Ccnb3^−/−^* oocytes that were left untreated, or 1 hour after strontium treatment in metaphase I (7h30 after entry into meiosis I). Chromosomes are stained with Hoechst (grey), centromeres with CREST (green). (B) Above, scheme of experimental setting to assess consequences of overexpression of cyclin B3 in wild-type CSF-arrested oocytes. Below, chromosome spreads of CSF-arrested oocytes injected with mRNA encoding WT or MRL mutant *M.m.* cyclin B3. (C) Time-lapse imaging of CSF-arrested oocytes injected as in (B) and incubated with SiR-DNA to visualize chromosomes. Quantification on the right shows the % of oocytes that exit CSF-arrest. (D) Control or Emi2 Morpholino (MO) knockdown *Ccnb3^−/−^* oocytes were used for time-lapse acquisitions using SiR-DNA. Scheme of Emi2 knockdown in *Ccnb3^−/−^* oocytes and quantification of oocytes that exit meiosis I are shown on the right. Arrow indicates anaphase I. All, error bars indicate means ± SD of the indicated number of oocytes from at least 3 independent experiments, indicated as dots. Scale bars: A/B: 10μm, C/D: 50μm See also Figure S1.

The key protein mediating CSF-arrest in mouse oocytes is Emi2 (Madgwick *et al*., 2006; Shoji *et al*., 2006). As we propose that *Ccnb3^−/−^* oocytes remain arrested in metaphase I because of precocious CSF-activity, knock-down of Emi2 should rescue progression through meiosis I in these oocytes. This was indeed the case: Injection of Emi2 morpholino oligonucleotides (MO) into prophase I arrested *Ccnb3^−/−^* mouse oocytes that were released to resume meiosis I, rescued chromosome segregation and polar body extrusion (Figure 1D). Consistent with the requirement of Emi2 for entry into meiosis II (Madgwick *et al*., 2006; Shoji *et al*., 2006), these oocytes then failed to proceed into the second meiotic division. Instead, they decondensed their DNA and entered interphase. The rescue of accomplishing the first meiotic division allowed us to conclude that untimely Emi2 activity in meiosis I accounts for the metaphase I arrest observed in *Ccnb3^−/−^*oocytes.

### Role of cyclin B3 is conserved in *Xenopus laevis* oocytes

The metaphase I arrest of *Ccnb3^−/−^*mouse oocytes can be rescued by expressing *Xenopus laevis* (*X.l.)* cyclin B3 (Karasu *et al*., 2019), indicating that cyclin B3’s role in vertebrate oocytes is evolutionarily conserved. This result opened up the exciting possibility that Xenopus oocytes can be used as a complementary model system for biochemical approaches not feasible in the mouse, to dissect the molecular function of cyclin B3. To this end, we first raised antibodies against a fragment (aa 1-150, Ab-B3^F^) and peptide (aa 6-26, Ab-B3^P^) of *X.l.* cyclin B3 and confirmed their specificity by Trim-Away. This technique exploits the E3 ligase and cytosolic antibody receptor TRIM21 to target endogenous proteins bound to antibodies for proteasomal degradation (Clift et al., 2017; Clift et al., 2018). Western blot (WB) analysis using Ab-B3^F^ confirmed that cyclin B3 was depleted in oocytes injected with TRIM21 mRNA and the peptide antibody Ab-B3^P^, but not control (Ctrl) Ab (Figure S2A). Unlike previously suggested based on WB data using a different antibody (Hochegger et al., 2001), cyclin B3 was expressed in prophase I Xenopus oocytes (Figure 2A). Upon progesterone (PG) treatment these oocytes resumed meiosis as shown by reduced SDS-PAGE mobility of the APC/C subunit Cdc27, phosphorylation of MAPK (indicating activation of the Mos/MAPK pathway which gradually increases as oocytes enter meiosis I and progress into meiosis II), and loss of inhibitory Cdk1 phosphorylation (a hallmark of Cdk1 activation) (Figures 2A and S1). As a macroscopic marker for successful meiotic resumption, we determined the percentage of oocytes displaying the characteristic white spot in their animal hemisphere (shown with an asterisk in Figure 2A), which is caused by pigment dispersal at the cell cortex mediated by germinal vesicle breakdown (GVBD). Cyclin B3 levels remained constant until about 1 hour post appearance of the white spot (4h post PG) and the subsequent decline in its levels coincided with a transient downshift of Cdc27 marking the meiosis I to meiosis II transition (Figures 2A and S1). Thus, cyclin B3 is expressed in *X.l.* female meiosis I, degraded at meiosis I exit and does not re-accumulate in meiosis II.

**Figure 2.**
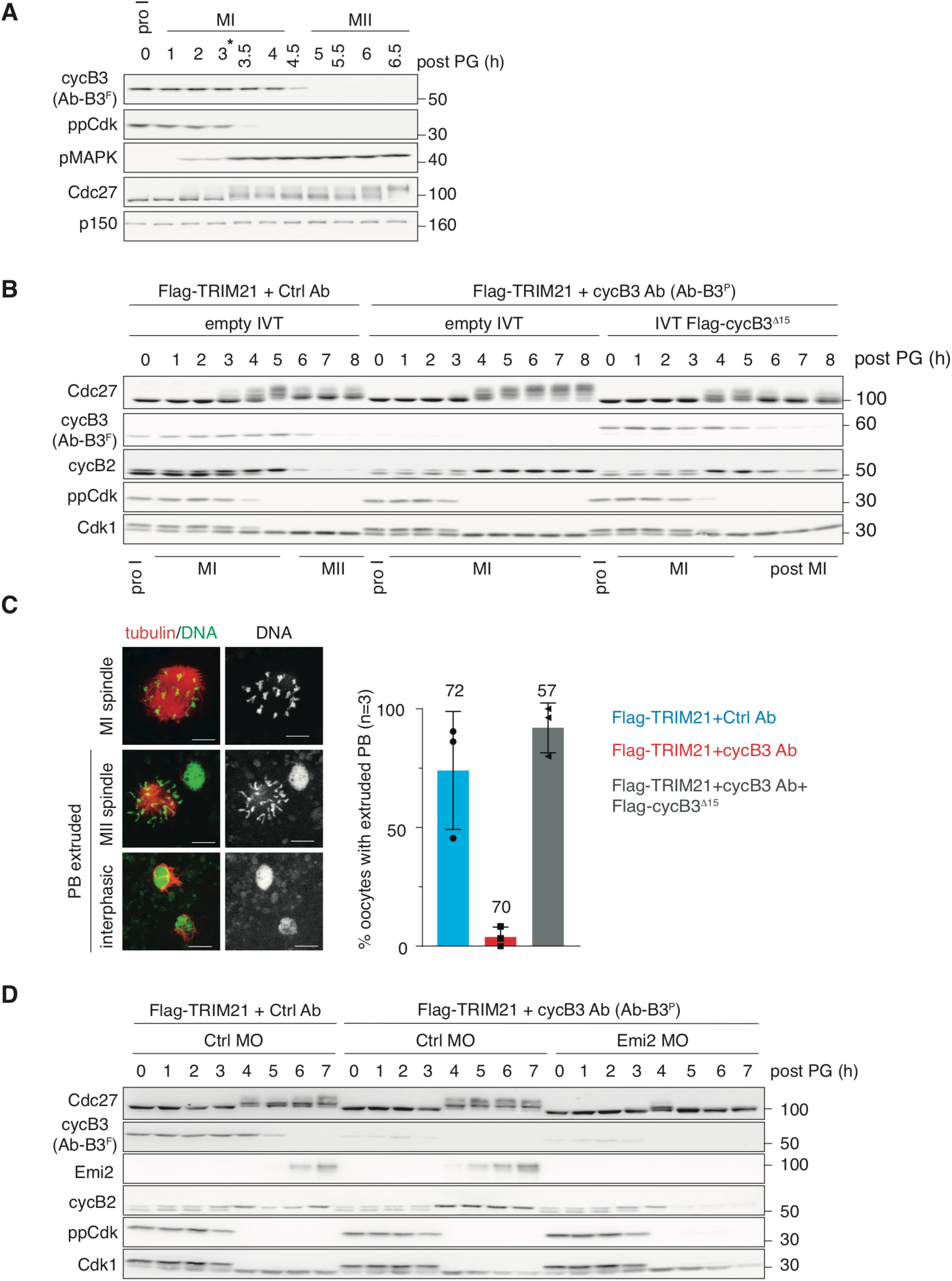
Loss of Xenopus cyclin B3 causes CSF-arrest in meiosis I. **(A)** Xenopus prophase I oocytes were treated with progesterone (PG) and at indicated time points immunoblotted for *X.l.* cyclin B3 (cycB3), Cdc27, inhibitory and activating phosphorylation of Cdk1 (ppCdk1) and MAPK (pMAPK), respectively. Asterisk marks the time when >80% of oocytes showed a white spot, indicative of oocytes that resumed the first meiotic division. p150 served as loading control. **(B)** Prophase I oocytes were injected with Flag-TRIM21 mRNA and control (Ctrl) or cyclin B3 peptide (Ab-B3^P^) antibody. Where indicated, IVT Flag-cyclin B3^1′15^ was co-injected for rescue experiments. 18 hours later, oocytes were treated with PG to induce meiotic resumption and at indicated time points analysed by WB. See also Figure S2B. **(C)** Oocytes treated as in B were microscopically analysed. Error bars indicate means ± SD of the indicated number of oocytes from 3 independent experiments (n), indicated as dots. Scale bars: 10μm **(D)** Prophase I oocytes were injected with Flag-TRIM21 mRNA and control Ab or Ab-B3^P^ together with control MO or Xenopus Emi2 MO. 18 hours later, oocytes were treated with PG to induce meiosis I and immunoblotted. See also Figure S2D.

Next, we used Trim-Away to assess the consequences upon loss of endogenous cyclin B3 in Xenopus oocytes. Prophase I oocytes were injected with Flag-TRIM21 mRNA and the peptide antibody Ab-B3^P^ or control Ab. At 18 hours post injection, cyclin B3 was efficiently depleted in prophase I oocytes injected with Ab-B3^P^ (Figures 2B and S2B). Control and cyclin B3-depleted oocytes efficiently resumed meiosis upon PG stimulation demonstrating that frog oocytes, like mouse oocytes, do not require cyclin B3 for efficient meiotic resumption (Karasu *et al*., 2019; Li *et al*., 2019) (Figure 2B). However, unlike control oocytes, cyclin B3-depleted *X.l.* oocytes were unable to exit meiosis I but remained arrested in metaphase I as judged by the presence of stable cyclin B2, persistently upshifted Cdc27, and failure of polar body extrusion (Figures 2B, C and S2B). Thus, the phenotype of Xenopus oocytes depleted of cyclin B3 mimics the one of *Ccnb3^−/−^* mouse oocytes. This phenotype was specific to cyclin B3 depletion because co-injection of *in-vitro* translated (IVT) Flag-tagged *X.l.* cyclin B3^Δ15^ rescued the meiosis I arrest (Figures 2B, 2C and S2B). The rescue construct lacks the first 15 aa and is therefore resistant to Trim-Away using Ab- B3^P^ (Figure S2C). Crucially, depletion of Emi2 by co-injecting Emi2 MO, allowed cyclin B3 depleted oocytes to exit meiosis I, as judged by the downshift of Cdc27 and cyclin B2 degradation (Figures 2D and S2D). As in the case of mouse oocytes, Xenopus Emi2 is required for entry into meiosis II (Ohe *et al*., 2007; Tang *et al*., 2008; Tung *et al*., 2007) and hence, these oocytes subsequently entered interphase as indicated by constantly downshifted Cdc27 and no re-accumulation of cyclin B2 (Figure 2D). Thus, in both Xenopus and mouse oocytes, loss of cyclin B3 results in a precocious, Emi2-mediated CSF-arrest in meiosis I suggesting that the function of cyclin B3 is evolutionarily conserved in vertebrate oocytes.

### Cyclin B3 targets Emi2 for degradation in a Polo-like kinase 1 (Plk1)-dependent manner

To understand how cyclin B3 suppresses the function of Emi2 as CSF, we utilized Xenopus CSF egg extract. CSF extract, prepared from metaphase II oocytes (Figure S1), is an ideal system for cell cycle analyses because it faithfully recapitulates the cell cycle processes occurring in intact oocytes and as an open system is easily amenable to experimental manipulations such as protein expression from mRNA. Furthermore, due to the absence of cyclin B3 in meiosis II (Figure 2A), ectopic cyclin B3 can be functionally analysed without interfering with the function of endogenous cyclin B3. CSF extract supplemented with mRNA encoding *X.l.* Flag-cyclin B3^WT^ was unable to maintain the CSF-arrest and exited meiosis II as indicated by cyclin B2 degradation (Figure 3A). As shown by lambda phosphatase (.) treatment, APC/C activation was mediated by Emi2 hyperphoshorylation followed by its degradation. Cyclin B3^MRL^, unable to bind Cdk1 (Figure S3A), did not destabilize Emi2 and accordingly the extract maintained the CSF-arrest (Figure 3A). Thus, expression of wild-type Xenopus cyclin B3 in CSF extract results in the identical phenotype observed upon expression of mouse cyclin B3 in mouse CSF-arrested oocytes (Figure 1C), i.e., override of the Emi2-mediated CSF-arrest.

**Figure 3.**
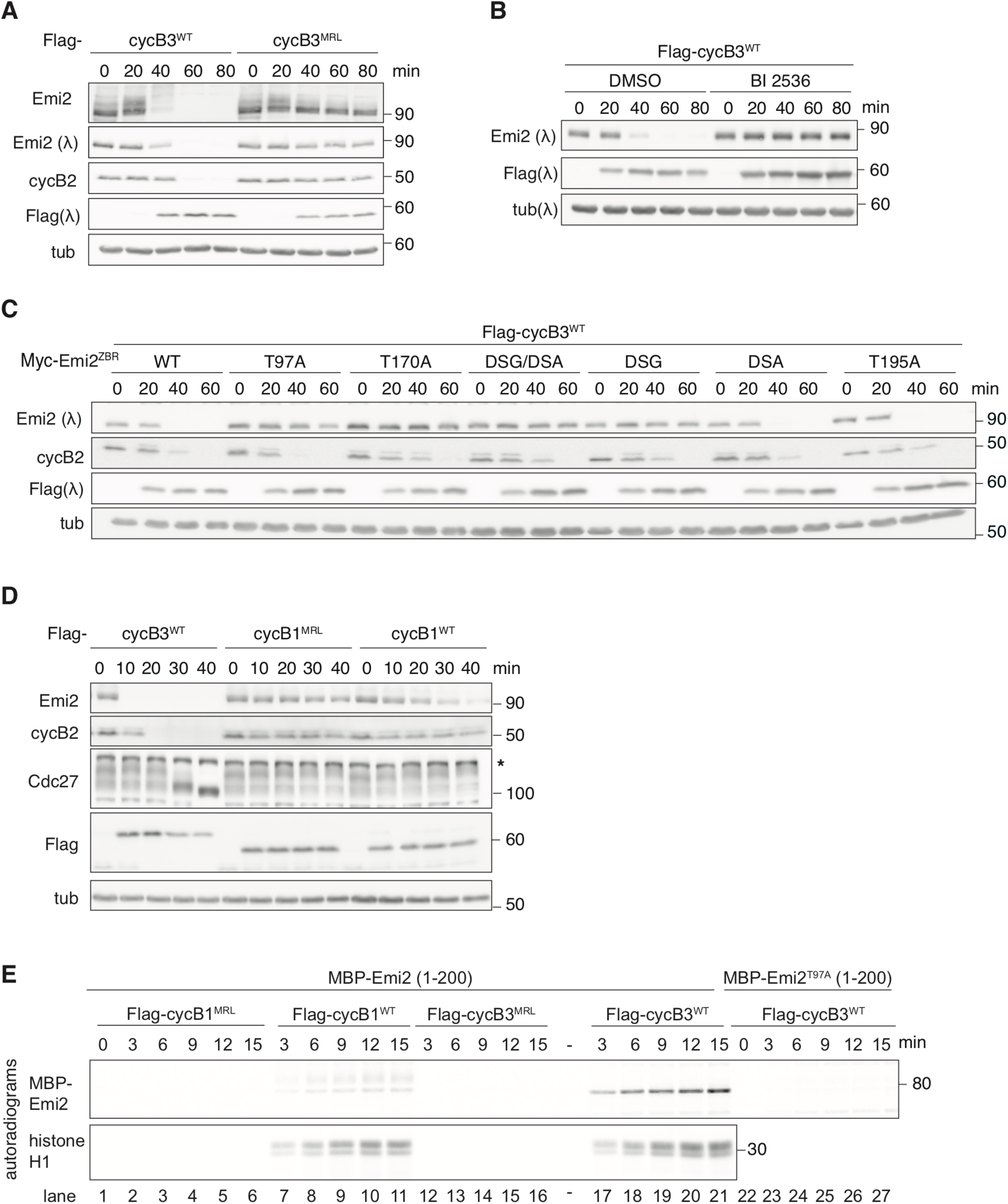
Xenopus cyclin B3 targets Emi2 for degradation in CSF extracts. **(A)** At the indicated time points after supplementing CSF extract with mRNA encoding Flag-tagged wildtype (WT) or MRL mutant (MRL) *X.l.* cyclin B3, extract samples were treated with lambda phosphatase (.) or not as indicated, and immunoblotted. Tubulin served as loading control. **(B)** CSF extract treated with 20μM BI2536 or DMSO was supplemented with *X.l.* Flag-cyclin B3^WT^ mRNA and samples were immunoblotted at indicated time points. **(C)** CSF extract was supplemented with IVT Myc-Emi2^ZBR^ variants and samples were immunoblotted at indicated time points after adding *X.l.* Flag-cyclin B3^WT^ mRNA. **(D)** IVT *X.l.* Flag-cyclin B1^WT^, -cyclin B1^MRL^, or -cyclin B3^WT^ was added to CSF extract and at indicated time points samples were immunoblotted. **(E)** *In vitro* kinase assay using Flag-tagged WT or MRL mutant of *X.l*. cyclin B1 (cycB1) or cyclin B3 (cycB3) expressed in CSF extracts followed by immunopurification. MBP/His-tagged Emi2 (aa 1-200) with S43, S73, and S157 mutated to A was used as substrate. Where indicated, the fragment contains an additional mutation at T97 (T97A). Autoradiogram and Coomassie-stained gels are shown. See also Figure S3D.

Previous studies showed that MPF activity (Maturation-promoting factor activity, corresponding to Cdk1 activity driving mitosis/meiosis) remains constant during Xenopus CSF-arrest despite ongoing cyclin B1 synthesis (Isoda et al., 2011; Wu et al., 2007b). At high MPF activity, Cdk1/cyclin B1 phosphorylates the N-terminus of Emi2 resulting in initial recruitment of Plk1, which then creates its own high affinity docking sites by phosphorylating Emi2 at T170 and T195 (Figure S3B). Subsequently, Plk1 targets Emi2 for degradation by SCF^β-TRCP^ through phosphorylation of two phosphodegrons (DSGX_3_S^38^ and DSAX_2_S^288^). The drop in Emi2 protein levels and the resulting transient APC/C activation reduces MPF activity, tipping the balance in favor of PP2A-B56, which dephosphorylates the inhibitory sites on Emi2 causing the re-stabilization of Emi2 (Inoue et al., 2007; Isoda *et al*., 2011; Nishiyama et al., 2007; Wu et al., 2007a; Wu *et al*., 2007b),. Hence, transitory phosphorylation of Emi2 controls APC/C activity to maintain MPF activity at constant levels while cyclin B1 synthesis continues.

Based on these data, we hypothesized that cyclin B3 might destabilize Emi2 via this pathway. Indeed, endogenous Emi2 remained stable when cyclin B3^WT^ was expressed in CSF extract supplemented with the Plk1 inhibitor BI2536 (Figure 3B). Next, we analysed the stability of ectopic IVT Myc-Emi2. Please note that we used Emi2 variants deficient in APC/C inhibition (ZBR) (Heim et al., 2018) to preclude that they interfere with cyclin B3 mediated CSF release. Mutation of the phosphodegron DSGX_3_S^38^ (DSG) strongly stabilized ectopic Emi2^ZBR^, while the DSAX_2_S^288^ (DSA) mutation had only a minor effect (Figure 3C). Consequently, the DSG/DSA double mutant was completely stable in CSF extract expressing *X.l.* cyclin B3^WT^. Mutation of T170 (T170A), but not of T195 (T195A), also resulted in the stabilization of Emi2^ZBR^ (Figure 3C). During CSF-arrest, Plk1 itself phosphorylates T170 and this depends on the Cdk1/cyclin B1 phosphorylation-dependent recruitment of Plk1 to Emi2’s N-terminus (Figure S3B) (Isoda *et al*., 2011). Crucially, Emi2^ZBR^ carrying non-phosphorylatable mutations at all four N-terminal Cdk1 sites (4 T/SP > 4A) was stable in the presence of cyclin B3^WT^ (Figure S3C). Of these sites, mutation of T97 (T97A), an evolutionarily highly conserved site (Figure S3B), was sufficient to stabilize Emi2^ZBR^ (Figures 3C and S3D). Thus, Plk1 activity as well as T97, T170 and the two phosphodegrons present in Xenopus Emi2 are critical for its degradation triggered by cyclin B3.

### Cyclin B3 and cyclin B1 in complex with Cdk1 differ in their *in vitro* substrate preference

Endogenous cyclin B1 cannot complement for the loss of cyclin B3 in mouse and frog oocytes. Furthermore, only addition of IVT WT *X.l.* cyclin B3, but not *X.l.* cyclin B1, induced rapid Emi2 degradation in CSF extracts, followed by CSF release (Figure 3D). Hence, our results indicate that Cdk1/cyclin B3 and Cdk1/cyclin B1 differ in their substrate specificity with only the former being able to efficiently phosphorylate Emi2 and thereby target it for fast degradation. To address this, we performed *in vitro* kinase assays using Flag-tagged cyclin B3 or cyclin B1 immunoprecipitated from CSF extract and a recombinant MBP-tagged Emi2 fragment (aa 1-200) as substrate. The respective MRL mutants served as negative controls. WB analyses of the IP fractions confirmed that Cdk1 was efficiently co-precipitated with the WT, but not MRL, versions of cyclin B3 and cyclin B1 (Figure S3D). Indeed, as shown by autoradiogram analyses WT cyclin B3, but not WT cyclin B1, efficiently phosphorylated Emi2^1-200^ (Figure 3E, upper panel, lanes 17-21 and 7-11). Note, that since T97 was most critical for the degradation of Emi2 (Fig. S3C), Emi2^1-200^ carried non-phosphorylatable mutations in the other three N-terminal Cdk1 sites to specifically detect T97 phosphorylation. We confirmed that Cdk1/cyclin B3 and Cdk1/cyclin B1 were equally active using histone H1 as generic Cdk1 substrate (Fig. 3E, lower panel). Notably, Emi2 with an additional mutation at T97 (T97A) was not detectably phosphorylated by WT Cdk1/cyclin B3 (lanes 22 – 27). From these data we concluded that cyclin B3 and cyclin B1 distinguish themselves in their substrate specificity in that Cdk1/cyclin B3, but not Cdk1/cyclin B1, can efficiently phosphorylate Emi2 on T97 targeting it for rapid degradation.

### Cyclin B3 destabilizes Emi2 in Xenopus meiosis I oocytes

Next, we validated our mechanistic insights *in vivo*, using intact Xenopus oocytes undergoing meiosis I. A key corollary of our hypothesis is that in the absence of cyclin B3 endogenous Emi2 should accumulate prematurely in meiosis I. To test this, we immunoblotted for Emi2 in control- or cyclin B3-depleted oocytes treated with PG. Since all samples were treated with lambda-phosphatase to correctly determine Emi2 levels and thus, all phosphorylation-dependent meiotic markers were lost, successful meiotic resumption was quantified using the appearance of the white spot as readout for GVBD (Figure 4A). In control-depleted oocytes, Emi2 did not appear until 5h post PG treatment. Note, that at 4h post PG all oocytes had already undergone GVBD. In contrast, in cyclin B3-depleted oocytes low levels of Emi2 were already detectable 3h after PG treatment, at a time in meiosis I when just ∼50% of oocytes had resumed meiosis I (Figure 4A). Premature Emi2 accumulation in early meiosis I was partially suppressed by co-injecting IVT Flag-cyclin B3^Δ15^ (Figure 4A). Notably, the effect of premature accumulation of Emi2 in the absence of cyclin B3 can also be observed in figure 2D. To further corroborate this finding, we analysed the stability of IVT Myc-Emi2^ZBR^ injected into cyclin B3-depleted and control oocytes stimulated with PG. Indeed, loss of cyclin B3 drastically stabilized Emi2^ZBR^ (Figure 4B). In line with our data from CSF-extract, mutation of T170, T97, or the two phosphodegrons also resulted in a significant stabilization of Emi2^ZBR^ in otherwise untreated oocytes undergoing meiosis I (Figure 4C). Altogether, we concluded that cyclin B3 prevents untimely CSF-arrest in Xenopus meiosis I oocytes by targeting Emi2 for degradation.

**Figure 4.**
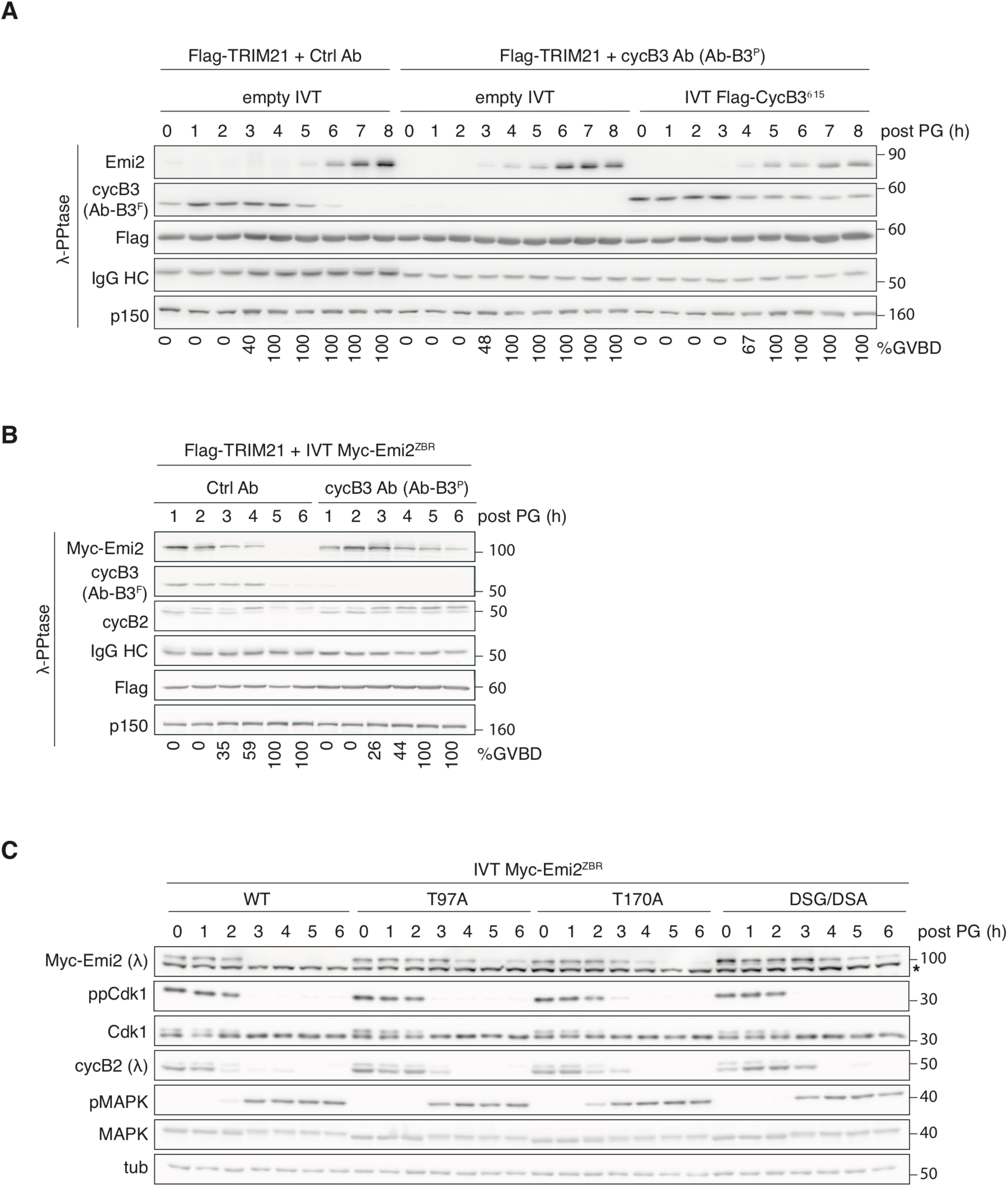
Xenopus oocytes lacking cyclin B3 prematurely accumulate Emi2. **(A)** Xenopus prophase I oocytes were injected with Flag-TRIM21 mRNA and Ctrl Ab or Ab-B3^P^ and empty or *X.l.* Flag-cyclin B3^Δ15^ IVT. 18 hours later, PG was added, and oocytes were immunoblotted at indicated time points. PG-induced meiotic resumption was confirmed by quantifying appearance of the maturation white spot, indicative of entry into prometaphase I. p150 serves as loading control. IgG HC: injected Ab heavy chain **(B)** Prophase I oocytes were injected with Flag-TRIM21 mRNA and Ctrl Ab or Ab-B3^P^ and 18 hours later with IVT Myc-Emi2^ZBR^. One hour later, oocytes were treated with PG and immunoblotted at indicated time points. **(C)** Prophase I oocytes were injected with indicated IVT Myc-Emi2^ZBR^ variants and at indicated time points after PG addition analysed by immunoblotting. Tubulin served as loading control.

### Cyclin B3 destabilizes Emi2 through a mechanism conserved from frog to mouse

Finally, we asked whether this molecular mechanism is conserved and cyclin B3 also targets mouse Emi2 for degradation. Since *X.l.* cyclin B3 (419 aa) is only one-third the size of mouse cyclin B3 (1396 aa), but still capable of rescuing *Ccnb3^−/−^* mouse oocytes (Karasu *et al*., 2019), we first tested whether *X.l.* cyclin B3 can induce the degradation of mouse Emi2 in Xenopus CSF extracts. Like in the case of Xenopus Emi2, we used a mouse Emi2 mutant (IVT Myc-Emi2^ZBR^) deficient in APC/C inhibition. Indeed, expression of *X.l.* cyclin B3^WT^, but not MRL mutant, destabilized mouse Emi2^ZBR^ in CSF extract (Figure 5A). Crucially, mutation of T86 (T86A), corresponding to T97 in *X.l.* Emi2 (Figure S3B), stabilized Emi2^ZBR^ (Figure 5A). Furthermore, both T152 (T170 in *X.l.* Emi2) and the only phosphodegron present in mouse Emi2 (DSGX_2_S^279^) were equally essential for its degradation, as deduced from the stability of the T152A and DSG mutants, respectively. These findings indicate that mouse and frog Emi2 indeed share the same mechanism of cyclin B3- dependent degradation.

**Fig 5.**
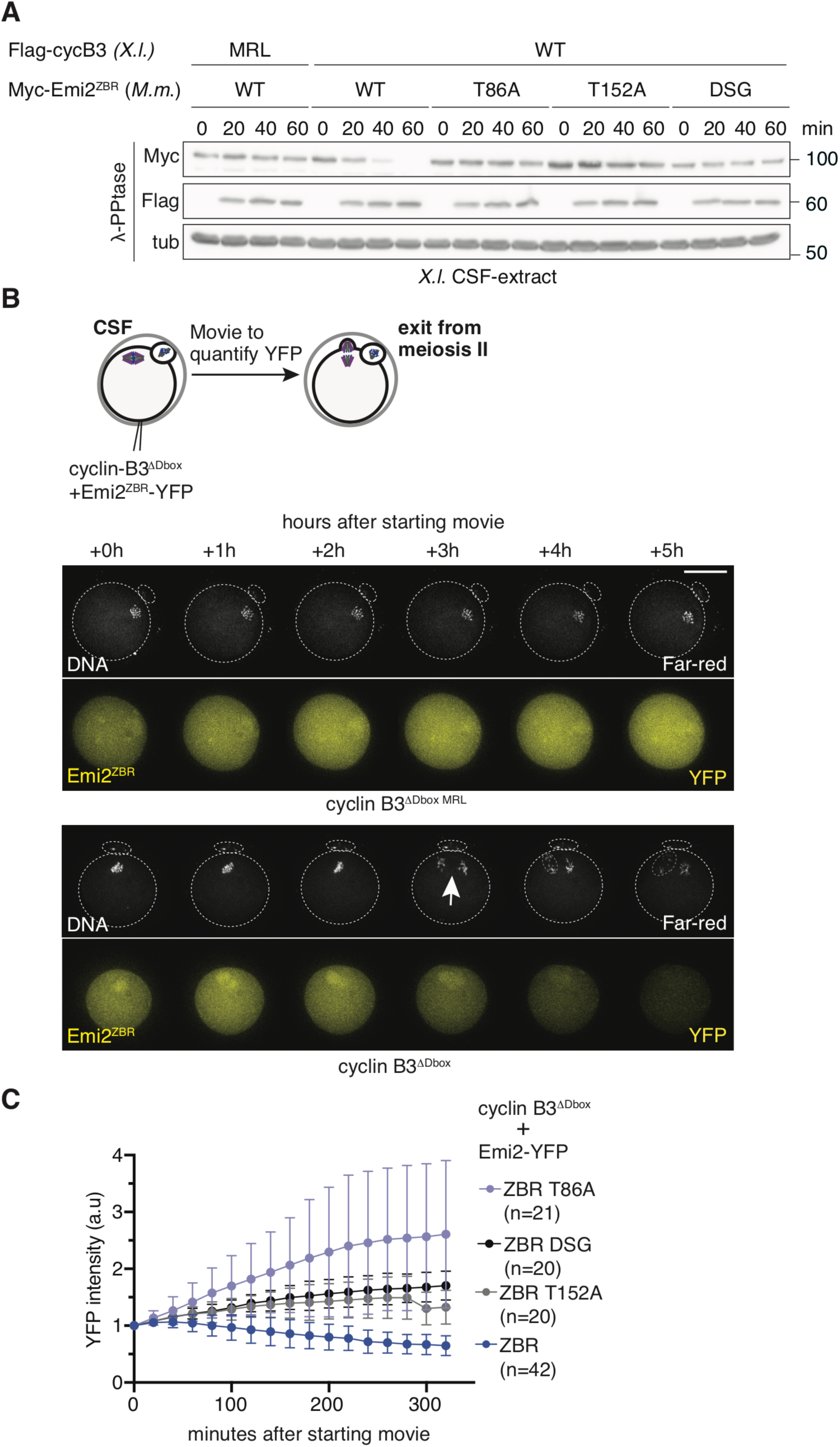
Cyclin B3 can target Emi2 for degradation in CSF-arrested mouse oocytes. (A) Xenopus CSF extract was supplemented with IVT *M.m.* Myc-Emi2^ZBR^ variants and at indicated time points after adding WT or MRL mutant *X.l.* Flag-cyclin B3 mRNA, samples were treated with lambda phosphatase (.) and immunoblotted. Tubulin serves as loading control. (B) Scheme illustrating the experimental setting and representative images showing time-lapse imaging of mouse WT CSF-oocytes injected with mRNA encoding Emi2^ZBR^-YFP and either cyclin B3^1′Dbox^ or cyclin B3^1′Dbox^ ^MRL^. SiR-DNA was used to visualize chromosomes. Anaphase II is indicated with an arrow. Scale bar: 50μm. (C) Quantifications of indicated Emi2^ZBR^-YFP mutants injected in CSF-arrested oocytes through acquisitions in the YFP channel (1z section) every 20 minutes with cyclin B3^1′Dbox^ expression. (B- C): n = number of oocytes from at least 3 independent experiments, error bars indicate means ± SD. See also Figure S4.

### Cyclin B3 targets Emi2 in CSF-arrested mouse oocytes

Next, we turned to mouse oocytes. However, due to the low number of oocytes obtained per mouse and low extract yield per oocyte, endogenous Emi2 can be only scarcely detected by WB analyses precluding the possibility to reliably quantify Emi2 levels. Therefore, we first analysed by live imaging the degradation of YFP-tagged Emi2^ZBR^ variants in CSF-arrested oocytes. Since endogenous cyclin B3 is not translated in mouse CSF oocytes (Han *et al*., 2017), we were able to investigate the effect of ectopic *M.m.* cyclin B3 on Emi2^ZBR^-YFP stability. Of note, we used stable *M.m.* cyclin B3 (ΔDbox) to ensure that protein levels were sufficient to efficiently target ectopic Emi2. Indeed, Emi2^ZBR^ was degraded in CSF oocytes co-injected with mRNA encoding *M.m.* cyclin B3^ΔDbox^, but not the corresponding MRL mutant (Figure 5B). Accordingly, oocytes expressing cyclin B3^ΔDbox^, but not the MRL variant, underwent anaphase II and extruded the second polar body. Exit from meiosis II indicates that endogenous Emi2 was also degraded upon ectopic expression of *M.m.* cyclin B3^ΔDbox^. Crucially, T86A Emi2^ZBR^-YFP was stable when co-expressed with *M.m.* cyclin B3^ΔDbox^ in CSF oocytes (Figure 5C). Additionally, mutation of T152 and the phosphodegron stabilized Emi2^ZBR^-YFP indicating that cyclin B3 in concert with Plk1 targets Emi2 for degradation. To further corroborate the function of Plk1 in Emi2 degradation, we investigated if CSF release induced by cyclin B3 expression depends on Plk1 activity. Indeed, Plk1 inhibition by BI2536 efficiently prevented CSF release induced by expression of ectopic *M.m.* cyclin B3^ΔDbox^ in mouse CSF oocytes (Figure S4B).

### Mouse cyclin B3 prevents CSF-arrest in meiosis I through an evolutionary conserved mechanism

Finally, we asked whether cyclin B3 controls Emi2 stability in its physiological context, i.e., mouse oocyte meiosis I. First, we analysed Emi2^ZBR^-YFP stability in WT oocytes progressing through meiosis I. We observed that Emi2^ZBR^-YFP was efficiently degraded in meiosis I, while the T86A variant was significantly stabilized (Figure 6A). To confirm that degradation of Emi2^ZBR^ during meiosis I in WT oocytes was mediated by cyclin B3, we analysed *Ccnb3^−/−^*oocytes expressing either WT or MRL mutant *M.m*. cyclin B3. Emi2^ZBR^-YFP was efficiently degraded in *Ccnb3^−/−^* oocytes expressing WT cyclin B3, but almost completely stable in MRL expressing oocytes (Figure 6B) demonstrating that Cdk1/cyclin B3 destabilizes Emi2 during meiosis I in mouse oocytes.

**Fig 6.**
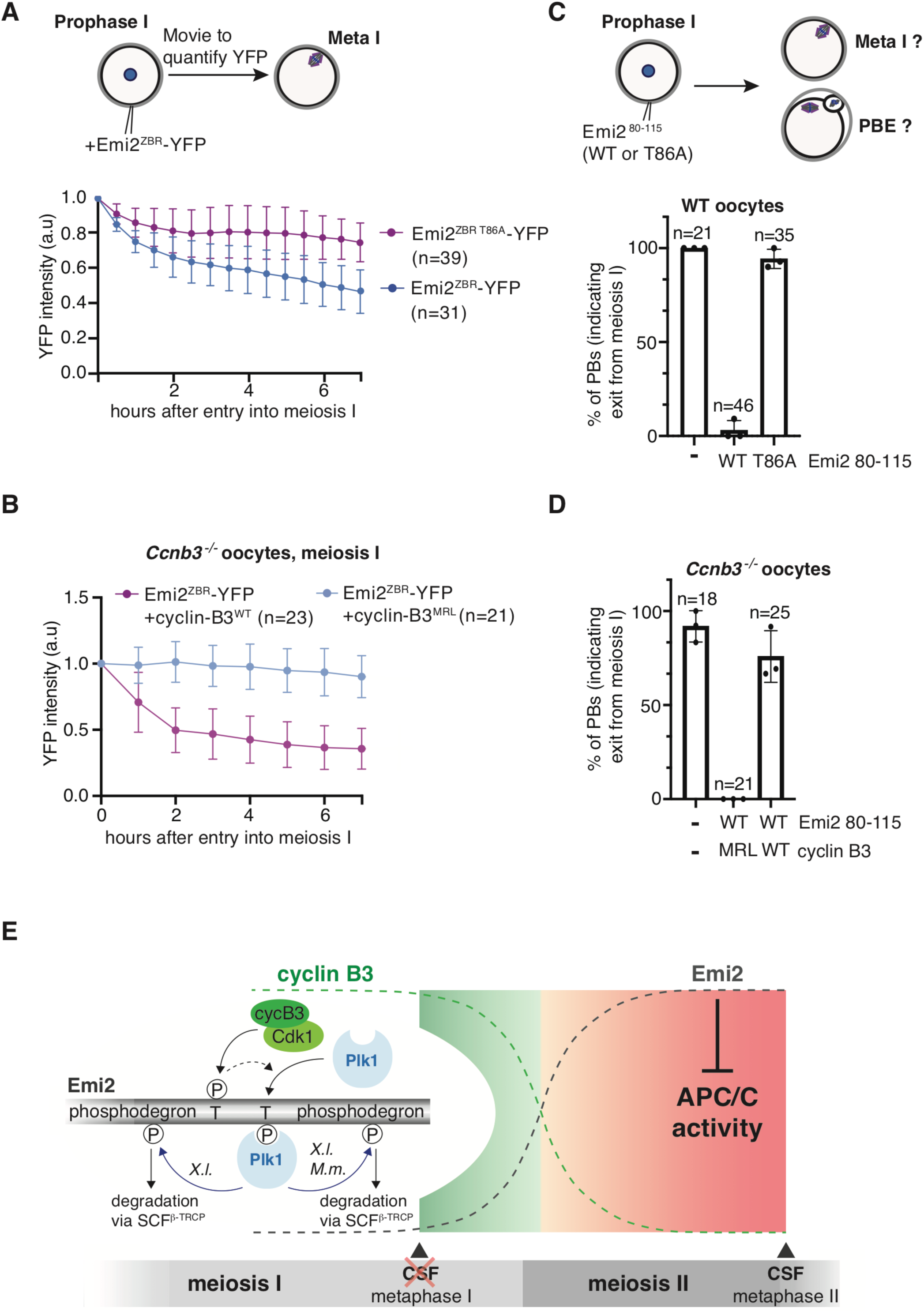
Cyclin B3 targeting of Emi2 for progression through meiosis I is conserved in mouse oocytes. (A) The experimental scheme above illustrates the analyses of Emi2^ZBR^-YFP stability in WT oocytes undergoing meiosis I. Prophase I-arrested oocytes were injected with Emi2^ZBR^-YFP or Emi2^ZBR^ ^T86A^-YFP and induced to resume meiosis I. YFP fluorescence was quantified every 30min (Error bars indicate means ± SD). (B) *Ccnb3^−/−^* prophase I oocytes were injected with mRNA encoding Emi2^ZBR^-YFP and either WT or MRL mutant *M.m.* cyclin B3. Following release into meiosis I, YFP signal was quantified hourly throughout meiosis I. (C) WT mouse prophase I oocytes were injected with WT or T86A Emi2^80-115^-RFP mRNA and following release, first polar body extrusion was quantified. (D) WT mouse prophase I oocytes were injected with WT Emi2^80-115^-RFP and either WT or MRL mutant *M.m.* cyclin B3 mRNA and quantified as in (F). (A-D): n = number of oocytes from at least 3 independent experiments, error bars indicate means ± SD See also Figure S5. (E) Model of how cyclin B3 prevents CSF-arrest in oocyte meiosis I. Degradation of Xenopus (*X.l.*) Emi2 is mediated via two phosphodegrons, while mouse (*M.m.*) Emi2 has only one phoshodegron. For further details, see text.

In mouse oocytes, expression of a small, 36 aa Emi2 fragment (aa 80-115) was shown to induce a metaphase I arrest by an unknown mechanism (Suzuki *et al*., 2010). Intriguingly, Emi2^80-115^ lacks the APC/C inhibitory domain (Figure S5A) and was only able to provoke an arrest when endogenous Emi2 was present (Suzuki *et al*., 2010). We therefore hypothesized that ectopic Emi2^80-115^, containing T86, outcompetes the phosphorylation of endogenous Emi2 by Cdk1/cyclin B3, resulting in stabilization of endogenous Emi2, untimely APC/C inhibition and hence, the observed metaphase I arrest. To test our hypothesis, we first expressed Emi2^80-115^-RFP in WT prophase I oocytes that were released into meiosis I. Indeed, these oocytes largely failed to exit meiosis I as indicated by the failure of first polar body extrusion (Figure 6C). Intriguingly, expression of T86A mutated Emi2^80-115^-RFP at comparable levels (Figure S5B) did not interfere with polar body extrusion and exit from meiosis I confirming our hypothesis that T86 phosphorylation of Emi2^80-115^ is critical for the observed dominant-negative effect of the fragment. If the metaphase I arrest was due to competition between Emi2^80-115^ and endogenous Emi2 for T86 phosphorylation by cyclin B3, increasing cyclin B3 levels should suppress the arrest. To test this, we used *Ccnb3^−/−^* oocytes ectopically expressing either WT or MRL mutant cyclin B3. Indeed, co-expression of WT, but not MRL mutant, cyclin B3 allowed to overcome the arrest caused by Emi2^80-115^ expression using PB extrusion as readout for exit from meiosis I (Figure 6D). In conclusion, also in mouse oocytes cyclin B3 in association with Cdk1 destabilizes Emi2 in meiosis I, a prerequisite for progression through meiosis I.

## Discussion

### Counting the meiotic divisions in oocytes to arrest at the correct time

In this study, we identify the mechanism that prevents vertebrate oocytes from installing a premature CSF-arrest already in meiosis I. Responsible for the timely arrest of mature oocytes at metaphase of meiosis II is Emi2, which directly inhibits the APC/C (Liu *et al*., 2006; Liu and Maller, 2005; Madgwick *et al*., 2006; Rauh *et al*., 2005; Schmidt *et al*., 2005; Shoji *et al*., 2006; Suzuki *et al*., 2010) and can only be found in vertebrates (Costache et al., 2014). During meiosis I, cyclin B3 prevents Emi2 from installing an untimely CSF-arrest by targeting it for degradation. The functional relationship between cyclin B3 and Emi2 we discovered provides the molecular explanation for the observation that *Ccnb3^−/−^* knock out mice are viable with fertile males, but infertile females (Bouftas and Wassmann, 2019; Karasu *et al*., 2019; Karasu and Keeney, 2019; Li *et al*., 2019). Male meiotic divisions come to an end before spermiogenesis is initiated; hence, once entry into the first meiotic division has taken place, the two meiotic divisions can run to completion without the necessity to establish a cell cycle arrest at any point (Lie *et al*., 2009). Therefore, the need for imposing an Emi2-mediated cell cycle arrest in female meiosis II comes at the cost of a need for cyclin B3, because it allows the oocyte to distinguish meiosis I from meiosis II.

Mitotic divisions do not require cyclin B3 and this is reflected by the fact that cyclin B3 knock-out mice are viable without any apparent phenotype, except female sterility (Bouftas and Wassmann, 2019; Karasu *et al*., 2019; Karasu and Keeney, 2019; Li *et al*., 2019). There are exceptions in non-vertebrates, for example the ascidian *Ciona intestinalis*, where cyclin B3 acts as a repressor of zygotic genome activation in embryos and contributes to the timing when the transcription machinery switches from maternal to zygotic mRNAs (Treen et al., 2018). In Drosophila, cyclin B3 was shown to promote anaphase onset during early embryonic mitotic divisions and female meiosis, but is dispensable later in development. Flies without zygotic cyclin B3 are viable but again, females and not males are infertile (Garrido et al., 2020; Jacobs et al., 1998; Yuan and O’Farrell, 2015). Notably, Drosophila lacks a true Emi2 orthologue and indeed it has been shown that Drosophila cyclin B3 directly activates the APC/C to promote anaphase in meiosis (Garrido *et al*., 2020).

### Emi2 destabilization in meiosis I

We show here that during female meiosis I, Cdk1/cyclin B3 in concert with Plk1 destabilizes Emi2 *via* an evolutionarily conserved mechanism to ensure that Emi2 levels are below the threshold critical for APC/C inhibition (Figure 6E). Based on our data, we propose a model where Emi2 degradation is initiated by the direct phosphorylation of T86/T97 (mouse/frog) by Cdk1/cyclin B3. Thereby, Cdk1/cyclin B3 serves as a priming kinase for Plk1 triggering the recruitment of Emi2 to Plk1, leading to phosphorylation of T152/T170 (mouse/frog) to create a previously identified docking site for Plk1 (Hansen *et al*., 2006; Jia et al., 2015). Plk1 then phosphorylates Emi2 at its phosphodegron(s) and thus generates recognition motifs for the E3 ligase SCF^β-TRCP^ resulting in efficient Emi2 degradation. To compare, in Xenopus oocytes at fertilization in meiosis II, Emi2 phosphorylation on T195 (another Plk1 docking site) by calcium/calmodulin dependent protein kinase II (CaMKII) primes Emi2 for Plk1, which then targets it for degradation by phosphorylating the phosphodegron (Liu and Maller, 2005; Rauh *et al*., 2005). Thus, Cdk1/cyclin B3 and CaMKII feed in the same degradation pathway depending on Plk1 resulting in fast and efficient Emi2 destruction, although with two completely different signaling inputs, i.e., activation of Cdk1/cyclin B3 during meiosis I versus calcium-triggered activation of CaMKII at fertilization.

Notably, expression of a short N-terminal fragment of mouse Emi2, lacking any APC/C inhibitory elements but comprising T86, causes a metaphase I arrest in mouse oocytes when endogenous Emi2 is present (Suzuki *et al*., 2010). Yet, the underlying mechanism remained elusive. Our data (Figures 6C and 6D) reveal that this metaphase I arrest is caused by titrating cyclin B3 away from endogenous Emi2 resulting in its stabilization and hence, APC/C inhibition.

### Expression of cyclin B3 exclusively in meiosis I

A central pillar of the mechanism underlying Emi2 accumulation in meiosis II is the concurrent absence of cyclin B3. As aforementioned, this meiosis I-exclusive expression of cyclin B3 allows the two meiotic divisions to be functionally distinct such that oocytes are only able to install CSF- arrest at metaphase of meiosis II, when the genome content has been halved. At exit from meiosis I, cyclin B3 – like the canonical B-type cyclins B1 and B2 – is targeted for degradation in a destruction-box and APC/C-dependent manner (Bouftas 2019). Importantly, however, and in contrast to cyclin B1 and B2, cyclin B3 does not re-accumulate in meiosis II (Figure 2A) (Han *et al*., 2017). A key question is therefore what makes cyclin B3 unique in that it is only expressed in meiosis I. In Xenopus oocytes, translation regulation of cyclin B3 is distinct from cyclin B1 (Pique et al., 2008): While cyclin B1 mRNA is translationally repressed in prophase I arrested oocytes and strongly activated upon progesterone stimulation, cyclin B3 mRNA, on the contrary, seems to be weekly repressed already during the arrest and not significantly activated upon progesterone treatment. Thus, translation of cyclin B3 mRNA seems to be controlled in a manner that ensures that cyclin B3 is present at sufficiently high levels already in early meiosis I and does not re-accumulate following its destruction at exit from meiosis I. For mouse oocytes, using loading of *Ccnb3* transcripts onto polysomes as readout for active translation, it has likewise been shown that cyclin B3 translation is strictly limited to meiosis I, while cyclin B1 and cyclin B2 mRNAs are efficiently loaded during both meiotic divisions (Han *et al*., 2017). Yet, the specific regulatory elements in cyclin B3 mRNA limiting its expression exclusively to meiosis I are unknown and further studies are needed to understand the underlying molecular mechanisms.

### Cyclin B3/Cdk1 substrate specificity for Emi2

Once oocytes progress into meiosis II, Emi2 becomes stabilized due to the absence of cyclin B3. During CSF-arrest of Xenopus oocytes, Cdk1/cyclin B1 phosphorylates Emi2 at multiple N-terminal sites that result in Emi2 inactivation as well as its destabilization *via* Plk1-mediated phosphorylation of the DSG/DSA phosphodegrons (Isoda *et al*., 2011; Wu *et al*., 2007a). Notably, T97 of *X.l.* Emi2, which we show here to be critical for Cdk1/cyclin B3 mediated degradation of Emi2, is part of the reported N-terminal Cdk1/cyclin B1 sites. This raises the important question of how oocytes can maintain the CSF-arrest with high Cdk1/cyclin B1 activity when at the same time Cdk1/cyclin B1 targets the essential CSF factor Emi2 for degradation.

As shown by our *in vitro* study (Figure 3E), T97 of *X.l.* Emi2 is a much better substrate of Cdk1 associated with cyclin B3 than with cyclin B1 demonstrating that cyclin B3 and cyclin B1 must distinguish themselves in their substrate specificity. While previous studies did not perform *in vitro* assays to assess T97 phosphorylation by Cdk1/cyclin B1, the *in vivo* phenotypes we observed argue against the idea of Emi2 being an optimal Cdk1/cyclin B1 substrate. First, the metaphase I arrest in mouse and frog oocytes upon loss of cyclin B3 was not rescued by endogenous cyclin B1 (this study and (Karasu *et al*., 2019)). Second, even ectopically expressed cyclin B1 failed to complement the lack of endogenous cyclin B3 (Karasu *et al*., 2019). Third, in CSF extract ectopically expressed cyclin B3 and cyclin B1 destabilize Emi2 with drastically different degradation kinetics (Figure 3D). While Emi2 is completely degraded within ten minutes after addition of IVT cyclin B3 WT, degradation of Emi2 in the presence of ectopic cyclin B1 WT is extremely slow and inefficient and does not result in CSF release. The slow and inefficient degradation of Emi2 by Cdk1/cyclin B1 matches its physiological function during CSF-arrest where it acts as part of a thermostat system that transiently destabilizes Emi2 to keep MPF activity constant despite ongoing cyclin B1 synthesis (Isoda *et al*., 2011). In sum, from these observations we concluded that differential substrate specificity of cyclin B3 and cyclin B1 is the cause why Emi2 is efficiently destabilized by cyclin B3 in meiosis I, while being stable in the presence of cyclin B1 in meiosis II.

In conclusion, our work elucidates a long-standing question, namely how vertebrate oocytes "count" the meiotic divisions, to arrest in meiosis II and not precociously, in meiosis I, for fertilization and successful embryo development. Our results demonstrate the essential and oocyte-specific role cyclin B3 occupies, by targeting the CSF-factor Emi2 for degradation through preferential substrate phosphorylation. Hence, cyclin B3 ensures timely oocyte arrest for fertilization. Loss of cyclin B3 in mice can result in triploid embryos upon fertilization or intracytoplasmic sperm injection (Chotiner et al., 2022; Li *et al*., 2019). Notably, mutations of *Ccnb3* in human oocytes have been associated with recurrent triploid pregnancies and miscarriage (Fatemi et al., 2020; Rezaei et al., 2021). Our work indicates that this phenotype is due to untimely CSF-arrest in oocytes devoid of functional cyclin B3. Future work will show how cyclin B3 substrate preference for Emi2 is brought about, and how cyclin B3 presence is restrained to meiosis I.

## Acknowledgements

We thank S. Keeney for comments on an earlier version of the manuscript and for *Ccnb3* knock- out mice, A. Dupré and S. Touati for discussion and comments, and M.E. Karasu for discussions at the start of the project. NB received a 1-year fellowship by the Fondation ARC pour la Recherche sur le Cancer. Work in the lab of TUM was funded by the German-Research-Foundation (MA 1559/10-1), and in the lab of KW by the Agence Nationale de la Recherche (ANR 16-CE92-0007- 01, ANR-19-CE13-0015), the Fondation de la Recherche Médicale (Equipe FRM DEQ 202103012574), a grant Emergence (OoCyclins, Paris Sorbonne Université), and core funding from the CNRS and Paris Sorbonne Université.

## Author contributions

Experiments on mouse oocytes were performed by NB with help from DC. FP performed initial experiments in Xenopus oocytes under the supervision of NB. Mouse husbandry and genotyping were done by DC, as well as expert technical help for all oocyte experiments. Experiments on Xenopus oocytes and with CSF egg extract were performed by LS, MH, MW, CK, JM, RD, HAA, PW, and AH. Cyclin B3 antibody was generated by MB. *In vitro* kinase assays were performed by RD. Figures were prepared by NB, LS, MH, RD, AH, TUM and KW. The manuscript was written by TUM and KW with input from all authors. Supervision, funding acquisition and project administration were done by TUM and KW.

## Supplemental figure legends

**Figure S1 (related to Figures 1 and 2).**
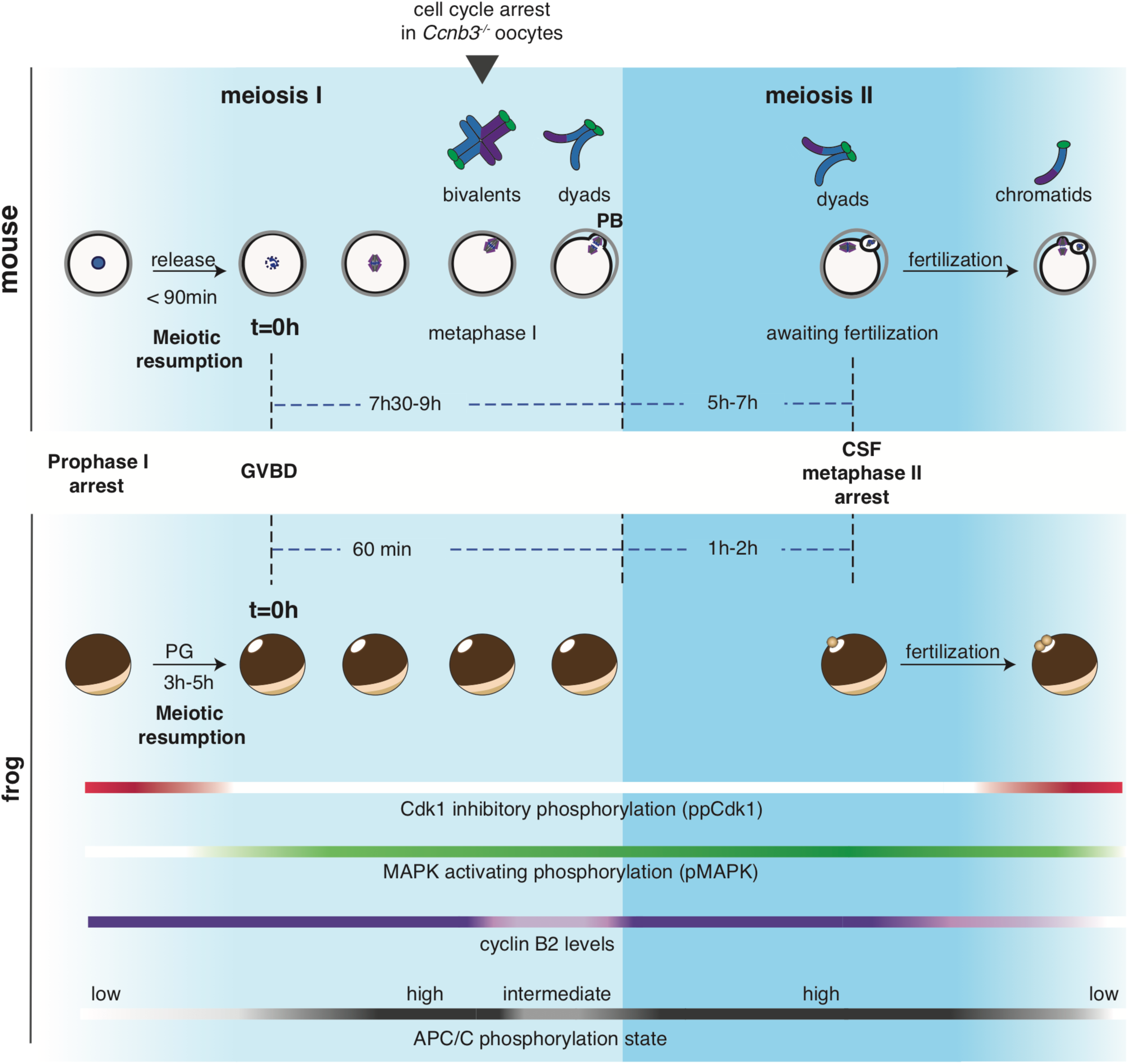
Schematic comparison of meiotic maturation in mouse and frog oocytes. **A**) Immature mouse and frog oocytes are arrested at prophase of meiosis I. Meiotic resumption is induced by washing oocytes in medium without dbcAmp (mouse) or by treating oocytes with progesterone (PG, frog). Germinal vesicle breakdown (GVBD) can directly be observed by light-microscopy in mouse oocytes, whereas the appearance of a white spot in the animal hemisphere of the oocytes serves as indirect readout for frog oocytes. Hours (h) and minutes (min) relative to GVBD are indicated. For mouse oocytes, subsequent meiotic phases can be monitored in real-time by live microscopy to follow chromosome movements and extrusion of the polar body (PB). Chromosome spreads allow further assessment of cell cycle stage (chromosome pairs (bivalents) in metaphase I, that segregate into paired sister chromatids (dyads) in anaphase I; in metaphase II, dyads that segregate into single sister chromatids in anaphase II.) Since opacity prevents live cell imaging of frog oocytes, Western blot analysis of indicated cell cycle regulators serves as readout for meiotic maturation in this organism. Loss of inhibitory phosphorylation of Cdk1 (ppCk1) and appearance of activating phosphorylation of MAPK (pMAPK) are used as markers for meiotic resumption. Retarded mobility of Cdc27 due to phosphorylation serves as readout for meiosis I. Polar body extrusion (PBE), cyclin B2 degradation, and downshift of APC/C mark exit from MI. Entry into meiosis II is accompanied by re-phosphorylation of APC/C. Mature oocytes await fertilization arrested at metaphase of meiosis II, also termed cytostatic factor- (CSF-) arrest. Spindles are in magenta, chromosomes in blue. Metaphase I arrest in *Ccnb3^−/−^* mouse oocytes is indicated by an arrowhead.

**Figure S2 (related to Figure 2).**
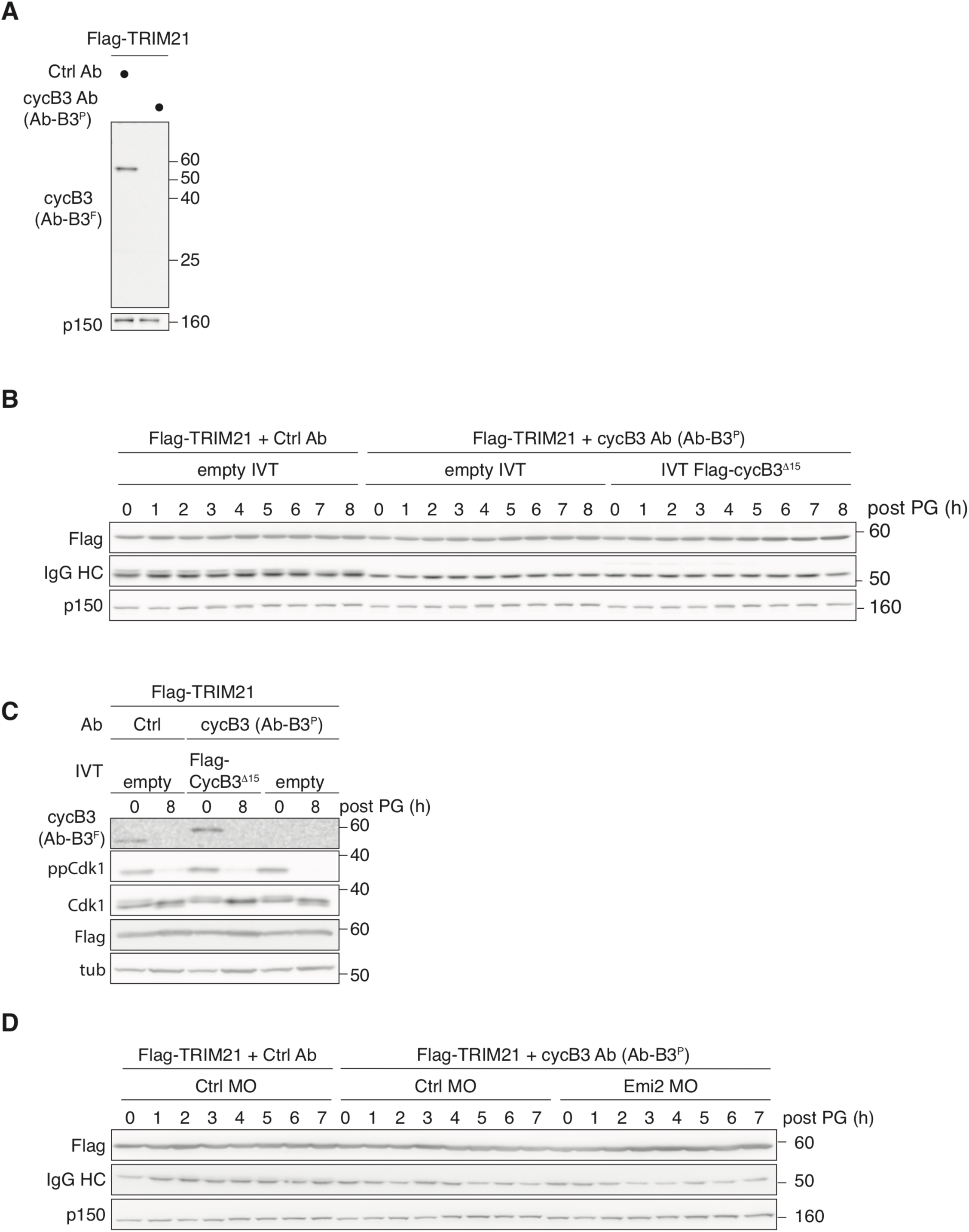
Validation of Xenopus cyclin B3 tools. **A**) Validation of cyclin B3 antibodies by TRIM-Away. Prophase I arrested oocytes were injected with Flag-TRIM21 mRNA and control (Ctrl) or cyclin B3 (cycB3) peptide Ab (aa 6-26, Ab-B3^P^). 23 hours post injection, oocytes were lysed and immunoblotted for cyclin B3 using the antibody raised against an N-terminal fragment of cyclin B3 (aa 1-150, Ab-B3^F^). p150 serves as loading control. **B**) Western blot (WB) showing expression of Flag-TRIM21, injected Ctrl or Ab-B3^P^ Ab (IgG HC: IgG heavy chain), and p150 as loading control for Figure 2B. **C**) Prophase I arrested oocytes were injected with Flag-TRIM21 mRNA, control (Ctrl) or cyclin B3 peptide Ab (Ab-B3^P^), and empty or *X.l*. *in vitro* translated (IVT) Flag-cyclin B3^Δ15^ and analysed by WB. Of note, cyclin B3^Δ15^ lacks the first 15 aa and is not recognized by the peptide antibody Ab-B3^P^, which was used to deplete endogenous cyclin B3 by TRIM-Away. Tubulin serves as loading control. **D**) WB showing expression of Flag-TRIM21, injected Ctrl or Ab-B3^P^ Ab (IgG HC: IgG heavy chain), and p150 as loading control for Figure 2D.

**Figure S3 (related to Figure 3).**
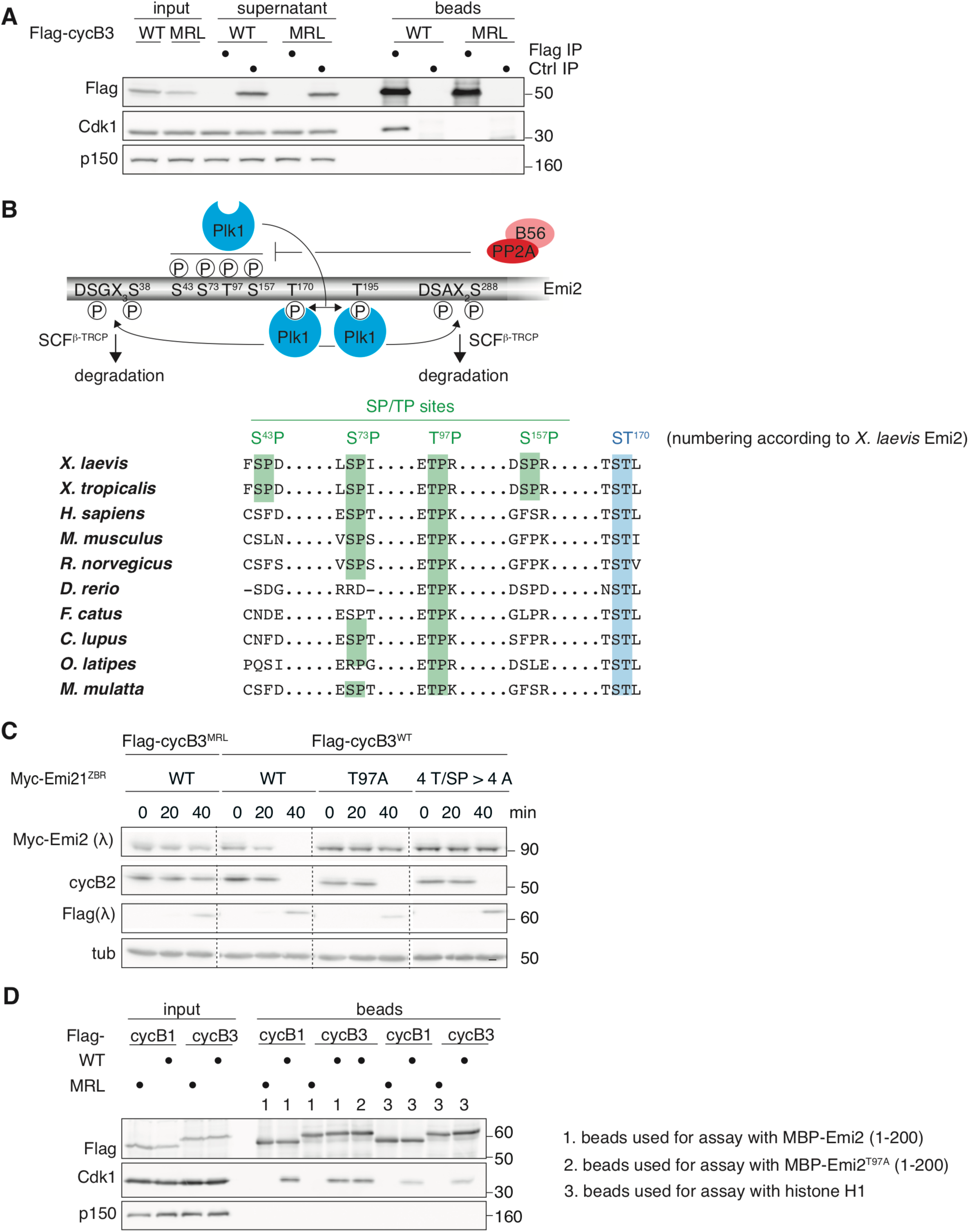
T97 is critical for cyclin B3-mediated Emi2 destabilization. **A**) Flag-tagged WT or MRL mutant *X.l.* cyclin B3 (cycB3) was expressed from mRNA in Xenopus CSF extract followed by immunoprecipitation using α-Flag or control (Ctrl) antibodies. Input, supernatant, and bead samples were analysed as indicated. p150 serves as loading control. **B**) Upper panel: scheme of Emi2 regulation during CSF-arrest to maintain constant MPF activity despite ongoing cyclin-B1 synthesis. See text for details. Lower panel: sequence alignment of Emi2 orthologues from different species. Numbering refers to *X.l*. Emi2. Green and blue sites refer to Cdk1 and Plk1 sites, respectively. **C**) Xenopus CSF egg extract containing IVT Myc-tagged Emi2^ZBR^ variants (WT: wildtype; T97A mutant; 4T/SP>4A: S43A, S73A, T97A and S157A) was supplemented with mRNA encoding Flag-tagged WT or MRL mutant *X.l*. cyclin B3 and analysed by immunoblotting. Where indicated, extract samples were treated with lambda phosphatase (.) prior to immunoblotting. Tubulin serves as loading control. **D**) CSF extract was supplemented with mRNA encoding Flag-tagged WT or MRL mutant *X.l.* cyclin B1 (cycB1) or cyclin B3 followed by anti-Flag immunoprecipitation. WB analyses of input samples and immunoprecipitates (beads), which were used for the *in vitro* kinase assay (Figure 3E) with the indicated MBP-Emi2 (aa 1-200) fragments or histone H1 as substrate. p150 serves as loading control.

**Figure S4 (related to Figure 5).**
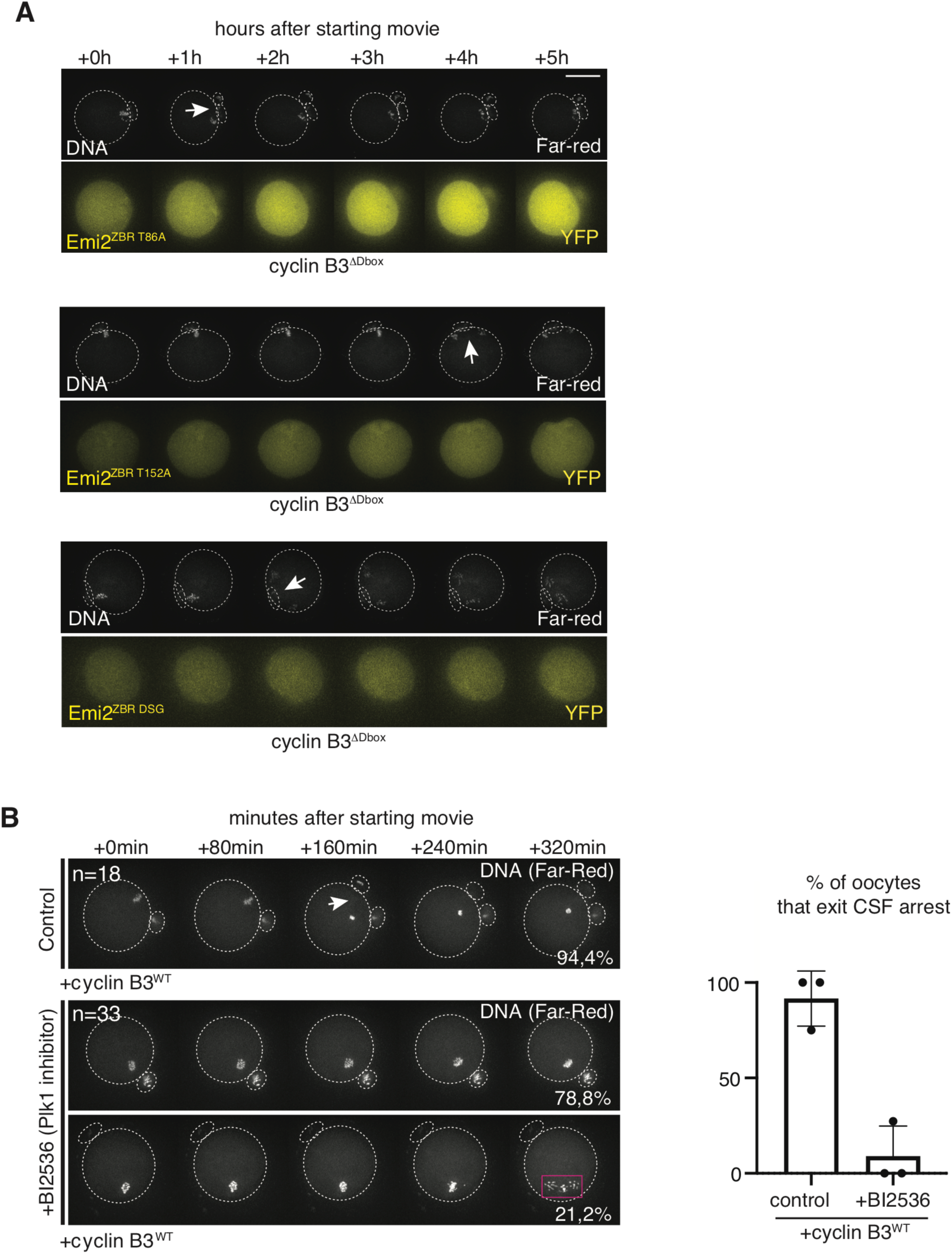
Live imaging of Cyclin B3-induced Emi2 degradation and CSF-release. **A**) mRNAs coding for the indicated Emi2^ZBR^-YFP mutants were co-injected with cyclin-B3^1′Dbox^ or cyclin-B3^1′Dbox^ ^MRL^ mRNAs in CSF-arrested mouse oocytes. Time-lapse acquisitions were done every 20 minutes. Selected images in the YFP channel (1z section) and overlays of the Far-red channel (15 z-sections, chromosomes) are shown, anaphase II is indicated with an arrow. Number of oocytes analysed is indicated in Figure 5C. Note: The 2nd PB was not always clearly visible, hence it is not necessarily indicated. **B**) CSF-arrested mouse oocytes expressing WT cyclin B3 were followed by time-lapse imaging in absence or presence of 50 nM of the Plk1 inhibitor BI2536. SiR-DNA was used to visualize chromosomes. Anaphase II in control oocytes is indicated with arrowhead, anaphase-like movements with a rectangle. Scale bar: 50μm. Oocytes that exit CSF-arrest or with perturbed segregation were quantified (Error bars indicate mean ± SD). n= number of oocytes from 3 independent experiments, means of each experiment indicated as dots.

**Figure S5 (related to Figure 6).**
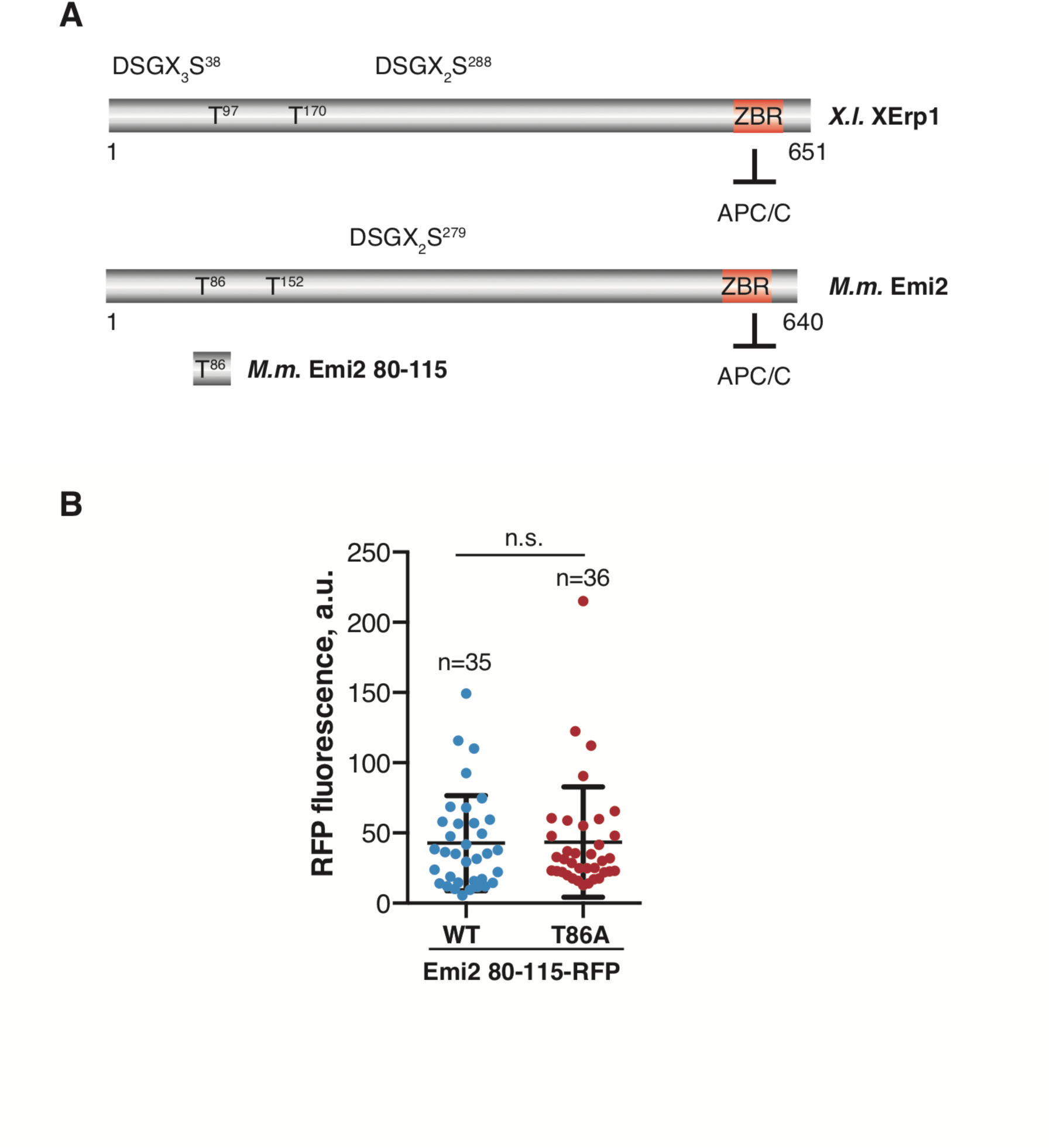
Emi2 fragment comprising T86 causes metaphase I arrest in mouse oocytes. **A**) Schematic representation of Mouse (*M.m*.) and Xenopus (*X.l.*) Emi2. Of note, the Emi2 fragment 80-115 lacks the APC/C inhibitory ZBR domain, but contains T86 (corresponding to T97 in frog), the Cdk1/cyclin B3 site, critical for cyclin B3-mediated degradation of Emi2. **B**) Qunatification of RFP-signal in oocytes that resumed meiosis expressing WT or T86A Emi2^80-115^- RFP. a.u.: arbitrary units, n indicates number of oocytes analysed, bar and whiskers are means and SD, n.s.: not significant, according to Mann-Whitney test.

## MATERIAL AND METHODS

### Animals

Mice were kept in a temperature, humidity and light controlled enriched environment with ad libitum access to food and water, at the animal facility of UMR7622 (authorization B75-05-13) and the IBPS (authorization A75-05-24), according to French regulations. Adult female CD-1 mice were obtained from Janvier, France, and Ccnb3−/− and Ccnb3+/− mice (C57BL/6/JRj background strain) were bred locally at the animal facilities. Genotyping was done as previously described (14).

Xenopus laevis frogs were bred and maintained under laboratory conditions at the animal research facility, University of Konstanz, and all procedures performed were approved by the Regional Commission, Freiburg, Germany (35-9185.81/G-17/121 and 35-9185.81/G-17/120).

### Mouse oocyte culture

To harvest mature prophase I oocytes, female mice were sacrificed by cervical dislocation by certified personnel at 8-16 weeks of age. Ovaries were dissected and oocytes recovered by mouth-pipetting follicles through a narrow glass pipet.Prophase I oocytes were cultured at 38° C in drops of M2 or M16 medium (homemade) supplemented with dbcAMP (dibutyryl cyclic AMP, Sigma Aldrich Merck, D0260) and covered with mineral oil (Sigma-Aldrich Merck, M8410). To release oocytes from prophase I arrest, they were washed three times in drops of M2 or M16 medium. Oocytes were resynchronized visually at GVBD (Germinal vesicle breakdown), and only oocytes undergoing GVBD within less than 90 minutes were used. *In vitro* culture conditions were controlled by verifying whether extrusion of the first polar body took place within 7,5 to 9 hours after GVBD in control oocytes. To induce release from CSF-arrest with Strontium, Ccnb3−/− oocytes at 7,5 hours after GVBD were put into M16 medium without CaCl2 (homemade) for 20- 30 minutes, and then transferred into activation medium (M16 medium without CaCl2 containing 100mM Strontium chloride, Sigma-Aldrich Merck 204463) for 1 hour. CSF-arrested Ccnb3+/− oocytes were used as controls for the activation procedure.

### Mouse oocyte microinjection

*M.m.* Cyclin-B3 pRN3 plasmids for in vitro transcription have been published (Karasu et al., 2019), Emi2 plasmids were PCR cloned into pRN3 for transcription from the T3 promoter. ZBR was mutated (C573A) with Pfu PCR mutagenesis kit. This plasmid was used as a template for the other Emi2 constructs, generated with Q5 site directed mutagenesis kit. Emi2 80-115 aa-RFP was generated with In-Fusion HD cloning kit (Ozyme). Primer sequences for cloning are available upon request.

mMessage machine kit (Invitrogen, AM1348) and RNAeasy purification columns (Qiagen, 74104) were used to obtain capped mRNA for microinjection into prophase I or CSF-arrested oocytes, using Eppendorf micromanipulators, Femto Jet, self-made microinjection pipettes (PN-31; Narishige) and constant flow settings. After injection, oocytes were left to recover in the incubator for 1-2 hours, unless otherwise specified. To study Emi2 stability in cyclin-B3-injected oocytes, Emi2 encoding mRNA was injected at least 30 minutes prior to cyclin-B3 mRNA. For MO knock- down of Emi2, Emi2 morpholinos (5’ ATTGCTTCCTGCTCTGTGGCTGGCT 3’) were injected into prophase I oocytes around 18 hours prior to release. Knock-down efficiency was verified by controlling for DNA decondensation and entry into interphase after first PB extrusion (Madgwick et al., 2006). For inhibition of Plk1, BI2536 was added at 50nM 1 hour before cyclin-B3 mRNA injection. Efficiency of Plk1 inhibition was controlled for by checking inhibition of anaphase I in control oocytes treated from GV onwards (Touati et al., 2015).

### Live imaging of mouse oocytes

Mouse oocytes were imaged under temperature-controlled conditions at 38°C on an inverted Zeiss Axiovert 200M microscope equipped with an EMCCD camera (Evolve 512, Photometrics), a Yokogawa CSU-X1 spinning disc, a nanopositioner MCL Nano-Drive, an MS-2000 automated stage (Applied Scientific Instrumentation) and piloted by Metamorph software, using a Plan-APO (63x/1.4 NA) oil objective (Zeiss). A final concentration of 1µM SirDNA (far-red DNA labeling probe, Spirochrome, SC007) was added to the culture medium 0,5-1 hour prior to imaging. At the indicated time points 11, 12 or 15 z-sections (3µm spacing) were acquired in the 640nm channel (Far-red) for SiRDNA imaging, and 1 z-section for 491nm channel for YFP, and the DIC image. Acquisitions for quantifications in Figure S5B were done on a Nikon Eclipse TE2000-E inverted microscope and PrecisExite High Power LED Fluorescence, a Prime sCMOS camera and a Plan APO (20x/0.75NA) objective. 1 z-section was acquired in the RFP channel for quantifications. Stills of movies were mounted with Fiji software, contrast and brightness was adjusted equally to all conditions being compared.

### Chromosome spreads

In short, zona pellucida-free mouse oocytes (tyrode’s acid treatment) were fixed in 0,65% - 1% paraformaldehyde, 0.15% Triton X-100 and 3mM DTT (Merck Sigma-Aldrich) at the indicated time points on microscopic slides (Chambon et al., 2013). Centromeres were stained with human CREST serum auto-immune antibody (Immunovision, HCT-100, at 1:50) and donkey anti-human Alexa Fluor 488 (709-546-149, Jackson Immuno Research, 1:200) secondary antibody, chromosomes with Hoechst 33342 (Invitrogen) at 50µg/ml. Slides were mounted with AF1 Citifluor mounting medium (Biovalley, AF1-100) or Vectashield (Eurobio H-1000), and examined with an inverted Zeiss Axiovert 200M spinning disc microscope as described for live imaging, using a 100X/1,4 NA oil objective. 6 z-sections with 0,4µm interval were taken. Images were mounted with Fiji software, contrast and brightness was adjusted equally to all conditions being compared.

### CSF extract preparation

3-14 days before the experiment, female Xenopus laevis frogs were injected with 20 U hCG subcutaneously into the dorsal lymph sack. 1 day (16-18 h) before the experiment, frogs were injected with 500U hCG to induce ovulation. The frogs were then placed into small tanks containing MMR (5mM Na-HEPES, 0.1mM EDTA, 0.1mM NaCl, 2mM KCl, 1mM MgCl2, 2mM CaCl2; pH 7.8). At the day of the experiment, laid eggs were collected, avoiding eggs with irregular pigmentation and apoptotic ones. Eggs of sufficient quality were washed with MMR and incubated in dejelling solution (2% cysteine, 0.1M KCl, 1mM MgCl2, 0.1mM CaCl2, pH 7.8) for up to 7 minutes, depending on oocyte surface state. Subsequently, eggs were washed with CSF-XB (100mM KCl, 1mM MgCl2, 0.1mM CaCl2, 50mM Sucrose, 10mM K-HEPES, 5mM EGTA, 1mM MgCl2, pH 7.7). Next, eggs were transferred to centrifuge tubes containing CSF-XB and Cytochalasin B (100µg/ml). The eggs were then compacted in a two-step centrifugation with the first step at 200 g for 1 minute and the second step at 650g for 1 minute. After removing residual buffer, eggs were lysed in a final centrifugation step at 16,500g for 10 minutes. From the resulting layers, the CSF-extract layer was taken with a syringe and mixed with Cytochalasin B (10µg/ml). From the resulting CSF-extract, a sample was taken and mixed with sperm nuclei to check for integrity of metaphase II arrest via the chromatin shape. Thereby, the DAPI stained chromatin appears as compacted fibers during the arrest and forms nuclei-like structures in interphase. A second sample was mixed with sperm nuclei and CaCl2 (0.6mM) to confirm the ability of the extract to release from metaphase II arrest. The extract was used at the day of preparation and kept on ice until used.

### Surgery and handling of prophase I arrested Xenopus laevis oocytes

To obtain oocytes arrested in prophase I, female Xenopus laevis frogs were anesthetized in Tricain solution (1g in 1L of MMR (5mM Na-HEPES, 0.1mM EDTA, 0.1mM NaCl, 2mM KCl, 1mM MgCl2, 2mM CaCl2; pH 7.8). To maintain a constant pH, 1g of NaHCO3 was added. The lack of reflexes was tested after 15 minutes to ensure proper anesthesia. Subsequently, skin and muscle were cut in a curved shape at either the left or right side of the abdomen. The ovary was partially removed by pulling with forceps and cutting with scissors. The removed parts were stored in MBS plus CaCl2 (88mM NaCl, 1mM KCl, 1mM MgSO4, 5mM HEPES, 2.5mM NaHCO3, 0.7mM CaCl2, pH 7.8). Muscle and skin were sewed separately and during recovery from anesthesia, the frog was placed in a separate tank with MMR, ensuring that the nose was above water. For microinjection, oocytes were then roughly separated in bundles of 15-20 oocytes and treated with 1mg/ml Liberase (Roche) to remove the surrounding tissue.

### Xenopus laevis oocyte injection and rescue experiments

For depletion of endogenous cyclin-B3, Trim21 mRNA (see section mRNA preparation for Xenopus oocytes) and anti-cyclin-B3 (Ab-B3P) antibody were injected into stage-VI oocytes. Here, 7ng of Trim21 mRNA and 20ng of antibody were used per oocyte. As control, unspecific rabbit IgG (20ng) were injected. For rescue experiments, IVT cyclin-B3715, not recognized by the antibody was co-injected (1.2 nl IVT per oocyte).

The oocytes were incubated overnight at 19°C in MBS medium (88mM NaCl, 1mM KCl, 1mM MgSO4, 5mM HEPES, 2.5mM NaHCO3, 0.7mM CaCl2, pH 7.8). Then, they were transferred to OR2 medium (82.5mM NaCl, 2.5mM KCl, 1mMCaCl2, 1mM MgCl2, 1mM Na2HPO4, 5mM HEPES, pH 7.8) containing 5µg/ml progesterone to induce meiotic resumption. Samples were taken every hour until GVBD spots appeared in the animal hemisphere of the oocyte, then every 30 minutes samples were taken. For Emi2 depletion via Morpholino oligonucleotides (MO) (GENE TOOLS, LLC) a mixture of Emi2 MO2 (AGATTTGCCATCTCTTGTTTCTT) and Emi2 MO3 (TGTGCCATCTCTTGTTTCTTTCTTC) was used. 9.2 pmol of each MO per oocyte were injected. As control the same amounts of inverse MO2 (TTCTTTGTTCTCTACCGTTTAGA) and inverse MO3 (CTTCTTTCTTTGTTCTCTACCGTGT) were injected.

### IF staining of Xenopus oocytes

For IF staining, oocytes were transferred to a fixation solution (100mM KCl, 3mM MgCl2, 10mM HEPES, 0.1% Triton, 0.1% glutaraldehyde, 3.7% formaldehyde, pH was adjusted to 7.8 before addition of Triton, glutaraldehyde and formaldehyde) and incubated overnight with slow shaking at 4°C. Then oocytes were bleached in 10% H2O2 in methanol for 20 h at room temperature exposed to light. Afterwards all incubation steps were done at 4°C in the dark (well-plates covered with aluminum foil). Oocytes were washed and blocked by incubating them 3 times for 1 h in AbDil buffer (0.1% Triton X-100, 2% BSA, 0.1% NaN3, in PBS). Next, oocytes were placed for 40h at 8°C in AbDil buffer containing FITC-labeled anti-tubulin antibody (1µg/ml final concentration) and Hoechst 33342 (1µg/ml final concentration). Oocytes were washed 4 times for 1 h by incubation in PBST on a shaker with gentle agitation. Finally, oocytes were mounted on glass slides in mounting solution (100mM KCl, 3mM MgCl2, 10mM K-HEPEs, pH 7.8, 0.1% Triton X-100, 50% glycerol).

### CSF time-course experiments

To analyze the stability of ectopic Emi2 in CSF extract, IVT reactions of the different Emi2 constructs were diluted 1:50 in CSF extract and incubated for 20 minutes at 20°C. Then, cyclin-B3 mRNA was added (either WT or MRL mutated) to CSF extract (final concentration: 90ng/µl) and samples were taken every 20 minutes up to 80 minutes. Where indicated, samples were treated with lambda phosphatase (NEB), for details see section “phosphatase treatment”. Finally, samples were mixed 1:10 with 1.5x Laemmli buffer (90mM Tris, 5% (w/v) SDS, 15% (w/v) glycerol, 12.5% (w/v) ß-Mercaptoethanol and Bromphenol blue) and heated for 10 minutes at 95°C.

### Phosphatase treatment

To facilitate the analysis of total protein levels, samples were treated with Lambda Protein Phosphatase (New England BioLabs). We found that for efficient dephosphorylation of CSF extract samples, conditions were slightly different from manufacturer’s instructions. Here, CSF samples were mixed 1:1 with phosphatase solution containing 1 volume PMP buffer, 1 volume MnCl2 buffer, 0.1 volume H2O and 0.1 volume lambda phosphatase. Then samples were incubated for 45 minutes at 30°C, and then mixed 1:5 with 1.5 x Laemmli buffer (90mM Tris, 5% (w/v) SDS, 15% (w/v) glycerol, 12.5% (w/v) ß-Mercaptoethanol and Bromphenol blue) and heated for 10 minutes at 95°C.

### SDS gels and Western Blot

For Western Blot samples, CSF extract was diluted 1:10 in 1.5x Laemmli buffer (90mM Tris, 5% (w/v) SDS, 15% (w/v) glycerol, 12.5% (w/v) ß-Mercaptoethanol and Bromphenol blue). Samples were heated at 95°C for 10 minutes and stored at -20°C until they were analyzed by immunoblotting. Per lane, 5µl samples were loaded corresponding to 0.5µl CSF extract.

Stage-VI oocytes were frozen in liquid nitrogen and stored at -80°C until lysis. 5µl of lysis buffer per oocyte (Mammalian Cell-PE LBTM Buffer (G-Biosciences) with 1x complete Protease inhibitor (Roche) were added, then scratched 10-15 times over a rack until a homogenous lysate formed. Then, the lysate was centrifuged 15 minutes at 20,000g at 4°C. From the resulting layers, the clear middle one was mixed 1:1 with 3x Laemmli buffer or underwent lambda phosphatase treatment with the same conditions as for CSF samples with the exception that here 5µl phosphatase solution were used for 18µl lysate. 5µl sample were loaded per lane to SDS gels, corresponding to half an oocyte.

For detection of Emi2, Cdc27 and cyclin-B3 8% SDS gels were used. For cyclin-B2, 10% SDS gels were used. The gels were run at 25mA for small gels (15 lanes) or 60mA for big gels (30 lanes). The proteins were then transferred to nitrocellulose membranes by wet blot. For small gels, the blot was run for 1h at 120V or 2h for big gels in wet blot buffer (25mM Tris, 0.19M glycine, 0.01% SDS and 20% methanol).

After blotting, membranes were blocked in 5% milk PBST or 3% milk TBST (for cyclin-B3) for 30 minutes at room temperature. The primary antibodies were incubated in the indicated dilutions overnight at 4°C. Before incubation of the HRP-coupled secondary antibodies (Dianova), the membranes were washed 3 times for 5 minutes in TBST (for cyclin-B3) or PBST (for all others). The secondary antibody solution was removed by washing 3 times for 5 minutes in TBST or PBST. For the detection of signals, membranes were incubated for 15sec in ECL solution (SuperSignal west Pico PLUS, Thermo Scientific), then the signal was detected with LAS-3000 (GE Healthcare Life Sciences).

### IVT preparation

Emi2 constructs were added as products from *in vitro* transcription and translation reactions. IVT reactions were done with the TNT® SP6 High-Yield Wheat Germ Protein Expression System (Promega) according to the manufacturer’s instructions. For stability tests of ectopic Emi2 constructs, IVTs were added at a 1:50 dilution to CSF extract.

### mRNA preparation for Xenopus oocytes

For expression of cyclin-B3 or Trim21, mRNA was added to CSF extract. The mRNA was prepared with the mMESSAGE mMACHINE® T7 Ultra Kit (Ambion) according to the manufacturer’s instructions. For testing Emi2 stability after expression of cyclin-B3 mRNA, 4.4µg of mRNA was added to 50µl CSF extract.

### IP of cyclin-B3 WT or MRL for kinase assays

Immunoprecipitation was performed with dynabeadsTM Protein G (Invitrogen). Beads were washed four times with PBST and once with PBS. Next, Flag antibody (Sigma-Aldrich) diluted in PBS was added and incubated overnight at 4°C rotating. Two-fold excess of beads were used for antibody binding compared to binding capacity described in the producer protocol. For example, for 100µl CSF-extract, 73.7µl beads were incubated with 9µg antibody. Before IP of Flag-tagged cyclin-B3, beads were washed three times with CSF-XB buffer.

For expression of Flag-cyclin-B3 WT or MRL, mRNA was added to CSF extract (final mRNA concentration: 40ng/µl) to validated CSF extract (see CSF extract preparation) and incubated for 30 minutes at 20°C. Through the addition of 100µM MG262 (Biomol GmbH) and IVT of a C-terminal fragment of Emi2 (aa 491-651, T545A, T551A; 1:20 diluted), maintenance of MII arrest was ensured. For immunoprecipitation, CSF extract was diluted 1:5 with CSF-XB containing 1x

PhosStop (Roche), 1x complete Protease inhibitor (Roche), and 0.375µM MG132 (Cayman). The IP took place for 90 minutes at 4°C rotating. The supernatant was discarded, and the beads were washed four times with PBS with 150mM NaCl and 0.5% Tween20 and two times with PB-buffer (1x PBS, 1% Glycerol, 20mM EGTA, 15µM MgCl, 100mM DTT, 1x complete, 1x PhosStop). Beads were maintained on ice until the start of the kinase assay.

### Kinase assay

For the kinase assay, purified fragments of Emi2 were diluted to a final concentration of 0.08mg/ml in PB-buffer (see section IP of cyclin-B3 WT or MRL for kinase assays) with 34.1µM ATP and 0.15µCi/µl [α-³³P]-ATP. For kinase assays using histone H1 as substrate, ATP and [α-³³P]-ATP were used at 44.9µM and 0.2µCi/µl, respectively. The zero-minute sample was taken previously to the assay start. The assay was started by adding recombinant Emi2 or histone H1 to the beads. Samples were taken at indicated time points. The assay took place at 25°C and mixed at 1,200 rpm. The kinase reaction was stopped by re-suspending the sample with 3x Laemmli buffer, followed by heat denaturation.

### Cloning Xenopus

The Xenopus laevis cyclin-B3 short isoform (Karasu et al., 2019) was cloned via PCR (Phusion polymerase kit, all primer sequences are available upon request) into pCS2 vectors containing either N-terminal Myc or Flag tag. The sequence is identical to NCBI NM_001085892.2. For the MRL construct, M190, R191, and L19A were mutated to alanine via PCR, the PCR reaction was digested with DpnI and subsequently transformed into *E. coli*. The different Emi2 constructs were cloned using a Morpholino-resistant Emi2 template by mutagenesis PCR with subsequent DpnI digestion. Emi2 constructs were cloned into PCS2 vectors containing either N-terminal Myc or Flag tag.

For time-course experiments, Emi2 constructs with mutations in the ZBR region (C583A) were used to prevent Emi2 from interfering with APC/C activity during the release from metaphase II arrest. Additional mutations were added to this construct with the oligos listed below, except for mutation of N-terminal XErp sites 4T/SP74A (S43A, S73A, T97A, S157A), where the DNA sequence containing the mutated sites was ordered from Thermofisher, cut with restriction enzymes EcoRV Hf (NEB) and FseI (NEB) and ligated with T4 ligase into pCS2 vector containing the residual Emi2 sequence. For DNA amplification, the constructs were transformed into *E. coli* Turbo (NEB) and DNA was recovered with Macherey-Nagel NucleoBond Xtra Midi kit.

Mouse Emi2 was cloned via PCR into a pCS2 Vector containing an N-terminal Myc-tag with oligos available upon request. The plasmid was used in a new PCR reaction generating the mutation of C573A (ZBR). This plasmid was used as template for PCR reactions introducing further mutations in the Emi2 sequence with Phusion polymerase.

### Quantifications of Emi2-YFP in mouse oocytes

For fluorescence intensity signal measurement, a circle of 60 × 60 pixels was placed on each oocyte, and another circle placed next to the oocyte for background values. For each oocyte, oocyte values were normalized after background substraction relative to the first value. Measurements were done with Fiji software using original, untreated acquisitions. GraphPad Prism 8 software was used for statistical analysis and at least 3 independent experiments were performed for each condition. The statistical tests applied are indicated in the corresponding figure legends.

